# MORPHIS (MORPHological Interpretable Signature) captures heterogeneous treatment- and aging-related responses of single cells

**DOI:** 10.64898/2026.03.24.713910

**Authors:** Freja Bohr, Athanasios Oikonomou, Emilie E. M. Nielsen, Georgios Konstantinidis, Janni S. Mortensen, Nektarios Tavernarakis, Hanne M. Nielsen, Nikos S. Hatzakis

**Author notes:** These authors contributed equally (Freja Bohr, Athanasios Oikonomou).

## Abstract

Cell morphology encodes changes in cytoskeletal and organelle organization during disease, treatment, and aging, yet is often assessed qualitatively or through poorly interpretable feature sets extracted from microscopy images. Here we introduce MORPHIS (MORPHological Interpretable Signature), a machine learning framework on explainable, analytically rich features paired with statistical methodologies for robust and interpretable quantification of single-cell morphological signature. MORPHIS extracts compact, interpretable feature signatures that capture both perturbation-specific response magnitude and heterogeneous cellular responses. It accurately distinguishes treatment-specific morphological signatures of eight mechanistically distinct membrane-active and intracellular-targeting compounds in Caco-2 and HeLa cells elucidating conserved and divergent phenotypic responses among compound-classes, as well as ultrastructural nuclear alterations upon aging of *C. elegans* and quantifies heterogeneous single-cell fractional responses. By remaining cell-type and perturbation agnostic, MORPHIS provides a generalizable framework for quantifying morphological signatures across diverse biological contexts including pharmacological treatment or aging.

## Introduction

Cell morphological characteristics like shape, size and other structural features(Keren et al. 2008; Prasad and Alizadeh 2019) are key descriptors of the physiological state of cells (e.g. cell division(Phillip et al. 2017), growth(Chambliss et al. 2013; Alizadeh et al. 2020; Grosser et al. 2021), metabolic activity, aging(Kamat et al. 2025), or migration(Adebowale et al. 2023)) and are therefore often used as a tool to identify cellular abnormalities in disease diagnostics(Singh and Lele 2022; Zhou et al. 2023; Sharma et al. 2025). Likewise, the specific functional cellular properties are often depicted by morphology(Lecuit and Lenne 2007; Prasad and Alizadeh 2019) (e.g. elongated axons in neurons or columnar structure of enterocytes). External stimuli such as chemical or mechanical stress can alter cell morphology, leading to changes in shape and structure (Wali et al. 2021; Tegtmeyer et al. 2024; Tang et al. 2024)). In a drug delivery perspective, drug treatment and cytotoxic stimuli are known to readily affect cellular morphology(Caliskan-Aydogan et al. 2024; Han et al. 2014; Kim et al. 2023), yet the mechanisms by which these changes occur remain poorly understood.

Traditionally, the common practice for assessing the effect of therapeutics (including the active pharmaceutical ingredient, excipients, and the drug delivery system) are primarily reliant on *in vitro* assays of cell viability, proliferation, and cytotoxicity (Adan et al. 2016). More specialized approaches also exist such as barrier integrity measurements of the trans-epithelial or endothelial electrical resistance (TEER)(Panou et al. 2023), reporter gene assays, or cell signaling and transporter assays. Phase-contrast and fluorescent imaging of fixed cells are often used in combination with the traditional ensemble assays but often rely purely on qualitative analysis(Panou et al. 2023; Palikaras et al. 2023). Most traditional assays report only ensemble-level quantitative readouts, which mask the inherent cell-to-cell heterogeneity within a biological population. Cellular responses are shaped by factors such as cell cycle state, stochastic gene expression, and local microenvironmental fluctuations(Snijder et al. 2009) leading to heterogeneous outcomes even among genetically identical cells. As a result, many perturbations elicit fractional responses, with subsets of cells exhibiting subtle morphological alterations while others display pronounced structural changes, giving rise to complex population-level phenotypes that are obscured by bulk measurements. Such fractional responses have been widely observed in contexts including apoptotic signaling(Spencer et al. 2009), antibiotic resistance(Balaban et al. 2004) and drug-induced senescence(Hernandez-Segura et al. 2018), where genetically identical cells adopt divergent morphological and functional states that are obscured by population-averaged measurements. Therefore, understanding the complex details of cellular morphology changes at the single-cell level likely will provide information of the underlying mechanism of any perturbation but also prove fundamental to understanding cellular function and aging mechanisms(Scipioni et al. 2025; Kounakis and Tavernarakis 2019; Palikaras et al. 2023).

Commonly, cell morphology is assessed qualitatively by interpretations from only a few representative sets of cell images(Bals et al. 2024; Lee et al. 2024). However, recent advancements in fluorescent imaging in combination with advanced computational analysis have allowed researchers to quantify the morphology of single cells(Lin et al. 2024; Tang et al. 2024; Janssen et al. 2022). Most morphological analyses rely on the extraction of descriptive features capturing the cellular morphology as used in CellProfiler(Carpenter et al. 2006) where the features are mathematically defined and describe, e.g. boundaries, area or intensity, normally relying on hundreds of generical features. While the extracted features can characterize the population, they do not, on their own, provide a platform for systematic comparisons across conditions or for identifying fractional responses at the single-cell level. Contemporary machine learning driven feature extraction utilizes deep learning to extract representations, without predefined features, designed for maximum classification accuracy often at the cost of interpretability for the user(De Vries et al. 2025; Phillip et al. 2021; Shen et al. 2025; Kleino et al. 2025; Lin et al. 2024; Salek et al. 2023; Sharma et al. 2025). This approach is especially used in cancer diagnostics with the development of unsupervised machine learning models to classify morphological differences in a cell population, identifying malignancy with higher speed and accuracy but with minimal focus on explainability(Piansaddhayanon et al. 2023; Kumar et al. 2024; Wang et al. 2026; Jost et al. 2025). Regardless of the methodological approach, however, extracting biologically interpretable mechanistic insights from these features remains challenging. Although such methods have been predominantly applied in cancer research, their use has not been fully extended to investigate the effects of drug formulations or other perturbations. Despite the advancements in quantitative image analysis combined with machine learning, contemporary research still lacks an end-to-end method that effectively captures biologically significant morphological changes and links them to specific treatments for extracting mechanistic insights.

Here, we combine fluorescent microscopy with quantitative image analysis(Bender et al. 2024; Malle et al. 2022; Bohr et al. 2023; Kæstel-Hansen et al. 2025; Gabriele et al. 2022; Caicedo et al. 2017; Thomsen et al. 2019) to capture single-cell morphological changes across multiple cellular compartments in intestinal epithelium monolayers and individually grown cells following treatment with eight test compounds, spanning from permeation enhancers (PE) and surfactants to anticancer drugs, inducing diverse morphological responses. We introduce MORPHIS, a machine learning framework that extracts treatment-specific morphological signatures by identifying the significant key morphological differences that contribute to the separation of classes allowing for easy interpretation of the factors contributing to the morphological change. The framework allows quantification of not only how a treatment alters morphology at a population level but also the magnitude and heterogeneity of responses at the single-cell level. We further extend MORPHIS to an *in vivo* model for studying aging, namely *C. elegans* demonstrating its applicability across experimental systems. These results establish MORPHIS as a versatile method for extracting interpretable insights at the single-cell level of cellular morphological alterations that operate robustly across treatments and cellular systems.

## Results

### MORPHIS: MORPHological Interpretable Signature

MORPHIS is a machine learning framework that operates on fluorescence images paired with segmentation masks of relevant cellular compartments (Fig. 1a) and can analyze two or more distinct perturbations between e.g. a control and test population or between different treatments. The framework works by first extracting a feature derived single-cell morphological signature from the combined fluorescent image and segmentation mask, consisting of 41 carefully selected features, capturing the distinct morphology of the cell, while maintaining a simple biological interpretation (Fig. 1b, see Supplementary Information Table 1 for a full list of features). These features cover quantification of shape, size, fluorescence intensity, texture and spatial compartment localization of two fluorescent channels (in this case nucleus and cytosol). MORPHIS employs a five-fold cross-validation strategy for classification with XGBoost, used to predict each cell’s condition based on the feature maps extracted (Fig. 1c). SHAP feature importance is applied to rank the most informative features, highlighting the key biological effects relevant to a specific treatment (Fig. 1c). Visualization of the single-cell prediction probabilities using UMAP (uniform manifold approximation and projection) encoding of the feature-space further reveals clusters of cellular responses, enabling exploration and quantification of both population-level treatment effects and heterogeneous, fractional responses at the single-cell level (Fig. 1d). MORPHIS generates outputs by summarizing complex morphological changes into interpretable treatment-specific signatures (Fig. 1e) by quantifying which cells respond to a specific treatment, how many and how strongly they respond, enabling a detailed assessment of effect at the single-cell level.

**Figure 1:**
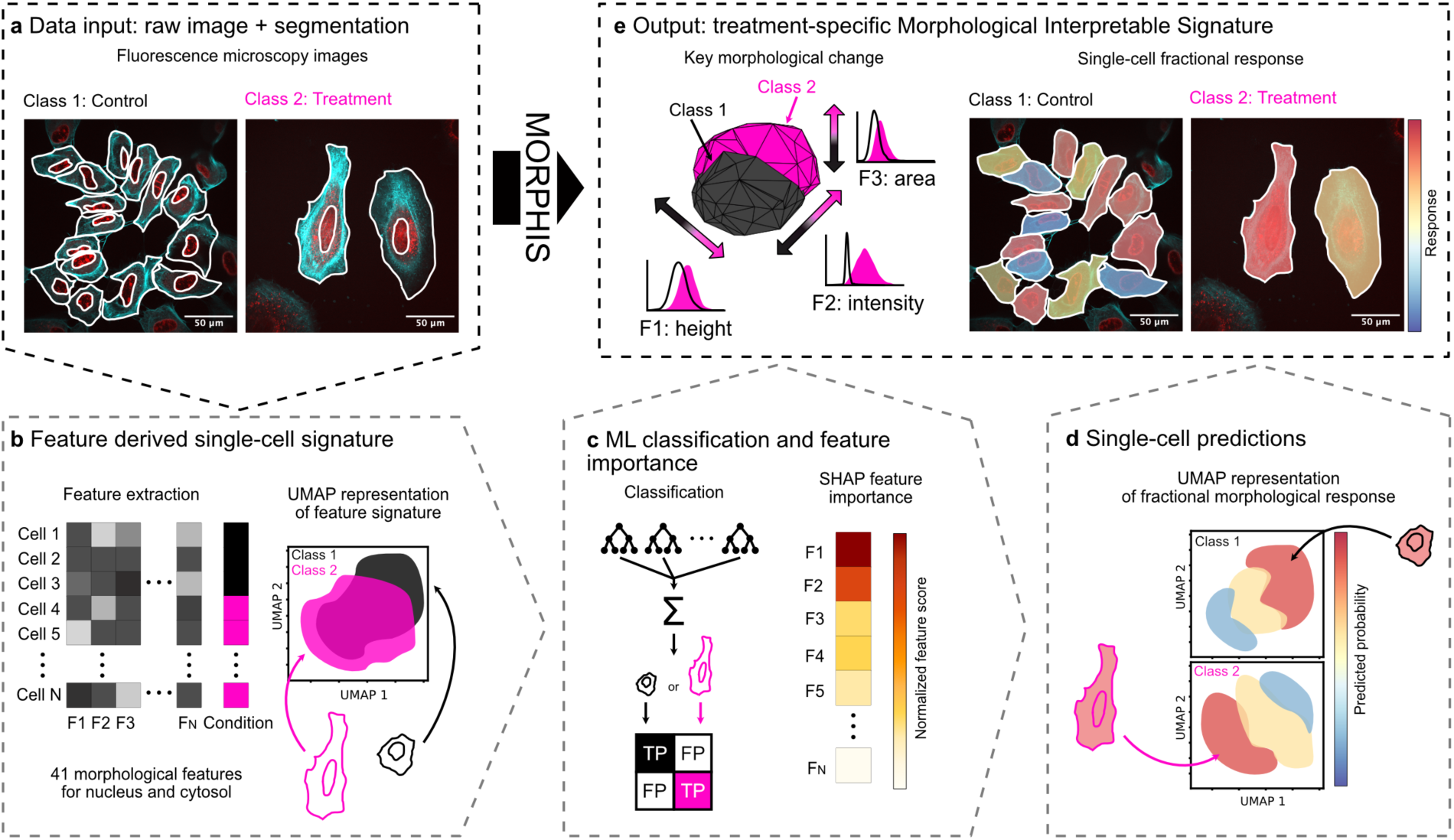
MORPHIS: An interpretable machine learning (ML) framework for quantifying treatment-specific single-cell morphological signatures and response heterogeneity. (a) Input: microscopy images of cells and segmentation masks derived from dual channel fluorescent markers that define single-cell nuclear and cytosolic compartments. (b) Feature level representation: For each cell, 41 biologically interpretable morphological features are extracted to generate a unique single-cell morphological signature. Features are categorized into groups: shape and size (e.g., area, circularity), intensity, and compartment organization (e.g., shannon entropy, homogeneity). These features form a single-cell morphological fingerprint used for downstream analysis. (c) Classification: XGBoost provides separability across conditions. SHAP feature importance: SHAP values rank morphological features according to their contributions providing key biological insight into what drives morphological variation. (d) Single-cell functional response: prediction probabilities provide a continuous measure for the single-cell fractional response. UMAP embeddings of feature-space reveal graded morphological transitions, enabling quantification of heterogeneous fractional responses rather than binary phenotypes at a single-cell level. (e) Output: treatment-specific signature of key morphological changes associated with treatment.

### Benchmarking MORPHIS across internal validation and competing methods

To assess the robustness and competitiveness of MORPHIS, we performed benchmarking analyses. First, multiple simple classifiers were evaluated in a binary classification task, distinguishing control cells from cells treated with the senescence-inducing anticancer drug doxorubicin (DOX) (Jost et al. 2025; Bisht et al. 2025; Wang et al. 2024; Sun et al. 2014). We compared the performance of a range of architectures (tree-based, support vector machines (SVM) and logistic classification architectures) (Fig. 2a, Supplementary Information Fig. 1). Model performance was evaluated using stratified five-fold cross-validation to obtain a robust estimate of generalization performance while preserving class proportions across folds. While performance varied modestly between classification models, accuracy remained consistently high above 78%, showing consistency of performance across models with the top-performing XGBoost model reaching 80.02±1.07%. Agreement in predictive performance across classifiers indicated that separability stems from underlying feature representation rather than model-specific effects. Visual inspection consistent with the SHAP feature importance rankings across classifiers revealed strong agreement in the top-ranked features (Fig. 2b, Supplementary Information Fig. 2) across models. In addition to confirming that the extracted morphological signatures are not dependent on classifier choice, this approach allows for the use of alternative classifiers when better suited to the task. UMAP embedding of the feature-space of four representative confidently predicted treated cells shown with the top five key features revealed coherent grouping (Fig. 2c), supporting the ability of the feature representation to capture graded morphological responses. Qualitative inspection further reveals broadly consistent treatment responses within each condition, while retaining subtle, yet detectable, differences in the biological feature-space.

**Figure 2:**
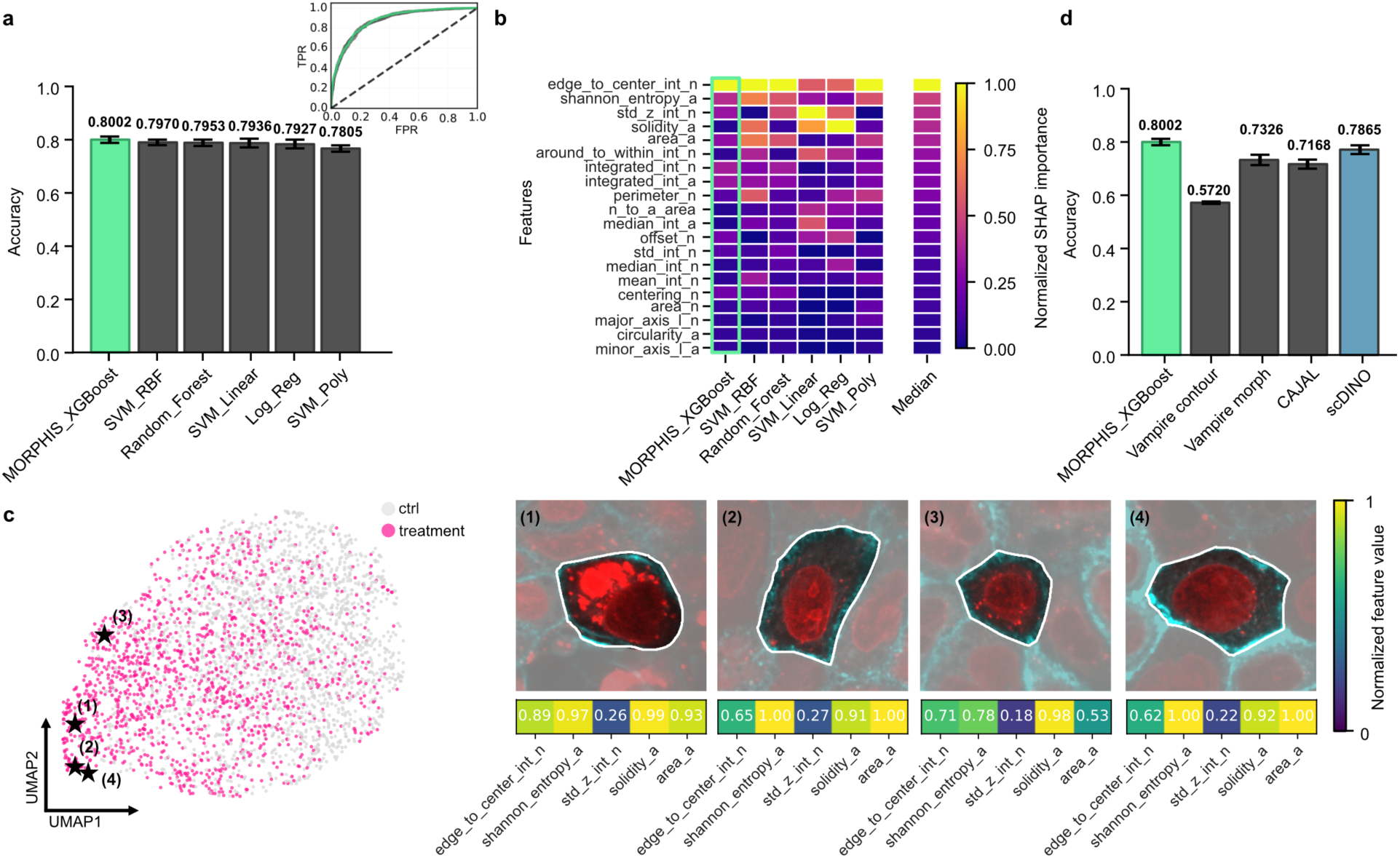
Benchmarking MORPHIS: internal validation and comparison to state-of-the-art methods. (a) Five-fold stratified validation accuracy for three support vector machines with different kernels (linear (SVM_Linear), polynomial (SVM_Poly) and radial basis (SVM_RBF)), random forest (Random_Forest), XGBoost (MORPHIS_XGBoost) and logistic regression (Log_Reg) for the specific binary classification task control (ctrl) vs. cells treated with the senescence-inducing drug doxorubicin (DOX). XGBoost achieved the highest performance accuracy (accuracy = 0.8002). Insert: ROC curves demonstrate identical model performance for the same binary task. (b) SHAP feature rankings for all tested classifiers show consistency in top important features regardless of architecture. (c) UMAP embeddings of the feature-space of the control and treated condition with four representative treated cells shown with the top five key features. Feature values are min-max normalized independently using pooled control and treated cells as the reference population. (d) Benchmarking of MORPHIS classification pipeline against state-of-the-art morphology analysis methods Vampire, CAJAL, and the deep learning method scDINO. MORPHIS is on par or achieves superior performance while retaining feature interpretability.

We next benchmarked the classification and feature extraction pipeline of MORPHIS against existing morphological analysis tools. The morphology of a cell entails spatial morphological components (e.g. size, shape, symmetry of signal), intensity (e.g. how and/or uniform is a measured biomarker per cell), or a combination of both. The unsupervised machine learning method Vampire(Phillip et al. 2021) (Visually Aided Morpho-Phenotyping Image Recognition) and neuronal cell shape characterization framework CAJAL(Govek et al. 2023) work explicitly on the masks of segmented cells extracting efficiently the spatial morphology of the segmented cells; accordingly, only the binary masks derived from image segmentation were used in these analyses. Because neither Vampire nor CAJAL include an integrated classification module, we extracted the features produced by each method and applied our classification pipeline on them. Vampire produced an accuracy of 57.20% for contour features and 73.26% for the six morphological features that Vampire automatically generates, while CAJAL reached 71.68% (Fig. 2d, Supplementary Information Fig. 3). These results indicate that the geometric shape of the cell is not enough to accurately describe the morphological effect.

**Figure 3:**
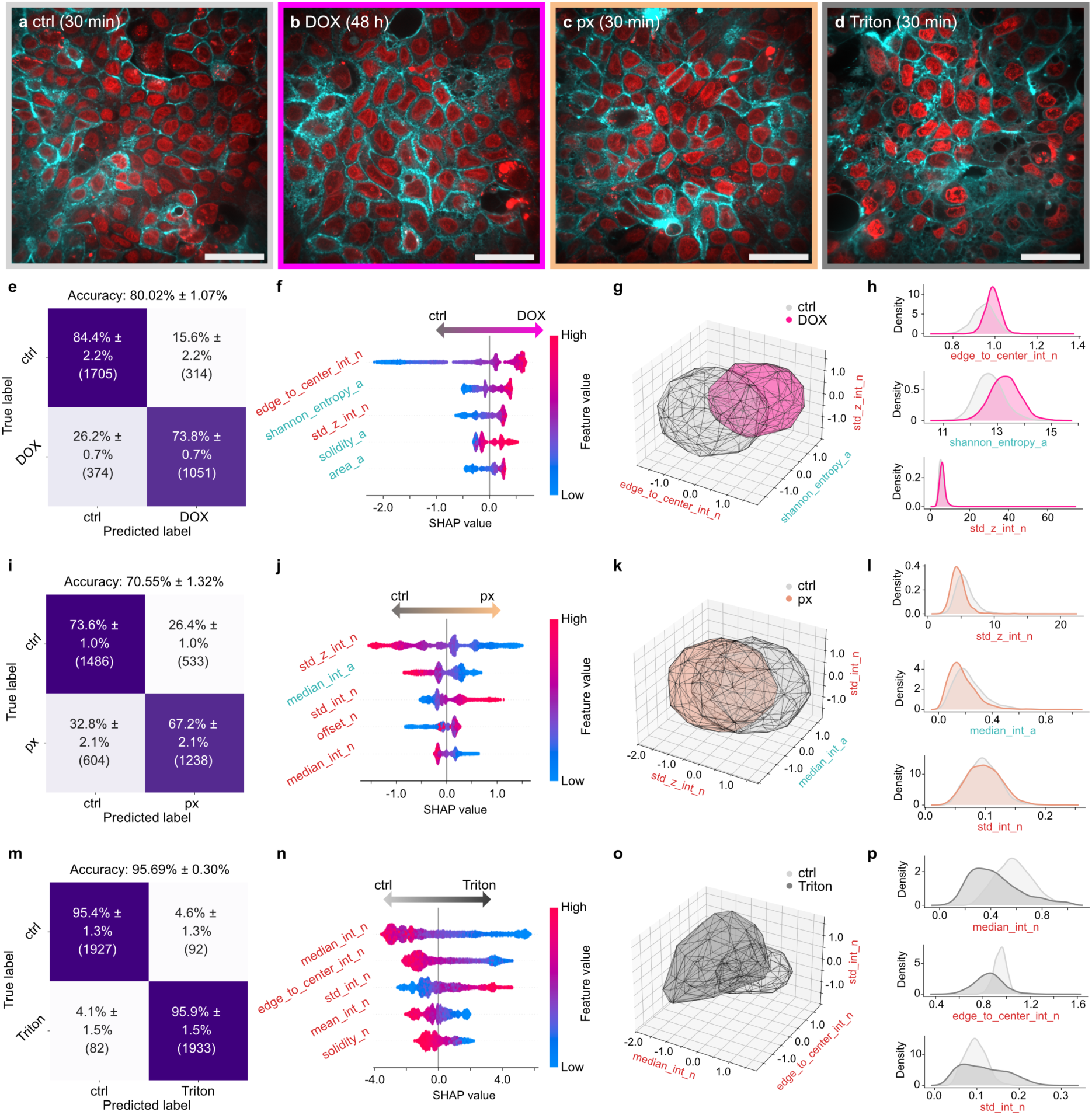
Treatment-specific morphological signatures revealed by feature importance of MORPHIS. MORPHIS was applied to classify Caco-2 monolayer cells under different conditions: (a) control (ctrl), (b) 3 µM doxorubicin (DOX) exposure for 48h, (c) 20 μM L-penetramax (px) exposure for 30 min and (d) 0.02% (w/v) Triton™ X-100 (Triton) exposure for 30 min. Images are representative average intensity projections (AIP) of z-stacks (0.5 μm between stacks) of non-fixed Caco-2 cell monolayers, with nucleus in red and actin in cyan. (e, i, m) Confusion matrices from five-fold cross-validated XGBoost classifiers with overall accuracy. (f, j, n) Beeswarm plots of the top five features ranked by SHAP values; feature names are colored by compartment: nucleus (red) or actin (cyan). The color bar denotes the range of numeric feature values. The arrow on top denotes the direction that a certain SHAP value will drive the prediction. (g, k, o) 3D convex hull plot of the top three SHAP-ranked features. (h, l, p) Kernel density estimation (KDE) plots of the top three features, with the highest-ranking feature on top. (scalebar = 50 μm, i = 4, n = 2, N = 3)

We further benchmarked against the state-of-the-art deep learning model scDINO(Hale et al. 2024; Pfaendler et al. 2023), which generates feature embeddings directly from cropped dual-channel cell images. For single-cell morphology quantification, we created a dual-channel object for each segmented cell in our dataset. For each cropped cell, scDINO creates a 364 long feature vector from the classification token that was passed on our classification pipeline achieving an accuracy of 78.65% (Fig. 2d, Supplementary Information Fig. 4a). Attention maps (Supplementary Information Fig. 4b) indicated broadly distributed focus across the entire cell area, consistent with global feature encoding thus providing information that is difficult to biologically interpret.

**Figure 4:**
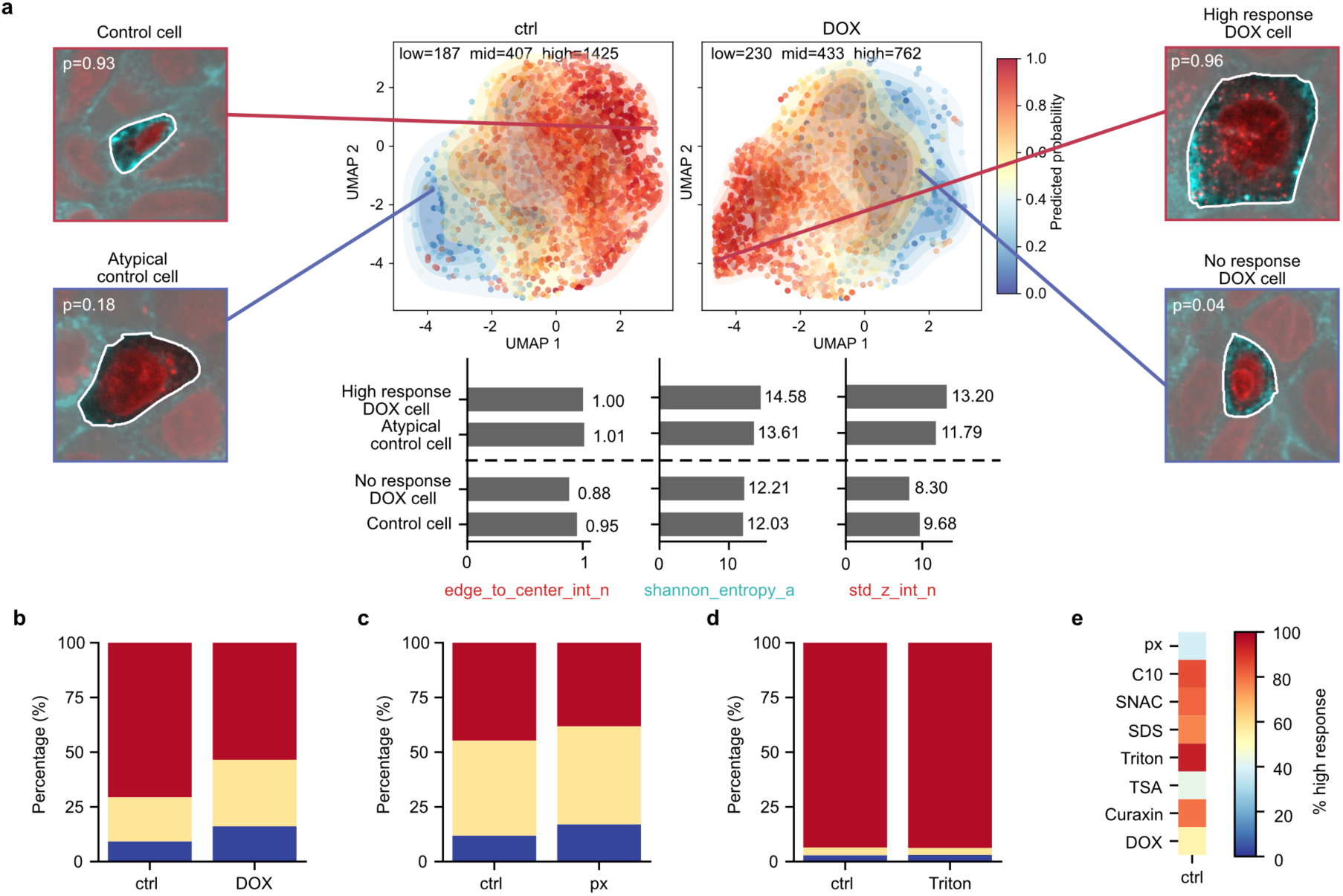
MORPHIS quantifies single-cell response heterogeneity and treatment-specific response distributions. (a) UMAP embedding of the feature-space for control (ctrl) and doxorubicin (DOX) treated Caco-2 monolayer cells. Points are color coded by the predicted class probability of belonging to a respective class XGBoost classifier. Four representative cells with low and high probabilities are shown. Underneath are the numeric top three feature values plotted for the representative cells. (b–d) Bar plots showing the distribution of single-cell predicted response probabilities for pairwise classifications of cells with low (p < 0.35, blue), mid (p = 0.35–0.7, yellow), and high (p ≥ 0.7, red) probabilities for: (b) control (ctrl) vs. doxorubicin (DOX) (c) control (ctrl) vs. L-penetramax (px) and (d) control (ctrl) vs. Triton™ X-100 (Triton). (e) Heatmap showing the fraction of high-response cells (p ≥ 0.7) for each treatment in the pairwise classification with control. (i = 4, n = 2, N = 3)

Collectively, this benchmarking demonstrates the robustness, generalizability, and competitiveness of MORPHIS. The consistent performance across classifiers and favorable comparisons with both classical shape-feature-based and state-of-the-art deep learning methods demonstrate that predictive power of MORPHIS arises from informative morphological features rather than from model-specific effects while at the same time enhancing biological interpretability of treatment-induced morphological changes.

### Morphological signatures of cellular responses to diverse treatments

To evaluate the versatility of MORPHIS, we conducted experiments and analyzed two different cell types representing distinctly different morphologies, namely the columnar enterocyte-like Caco-2 cells in a confluent monolayer and non-confluent epithelial HeLa cells. Eight treatments spanning PEs, surfactants, and anticancer drugs were selected as they were expected to induce diverse morphological changes. To ensure compatibility across treatments, the exposure concentration was tuned to the one resulting in ∼80% cell viability relative to buffer controls, as assessed by MTS-PMS assay (Supplementary Information Fig. 5-6 & Supplementary Information Table 2). For PEs and surfactants, the exposure time was 30 min, while for the anticancer drugs it was 48h.

**Figure 5:**
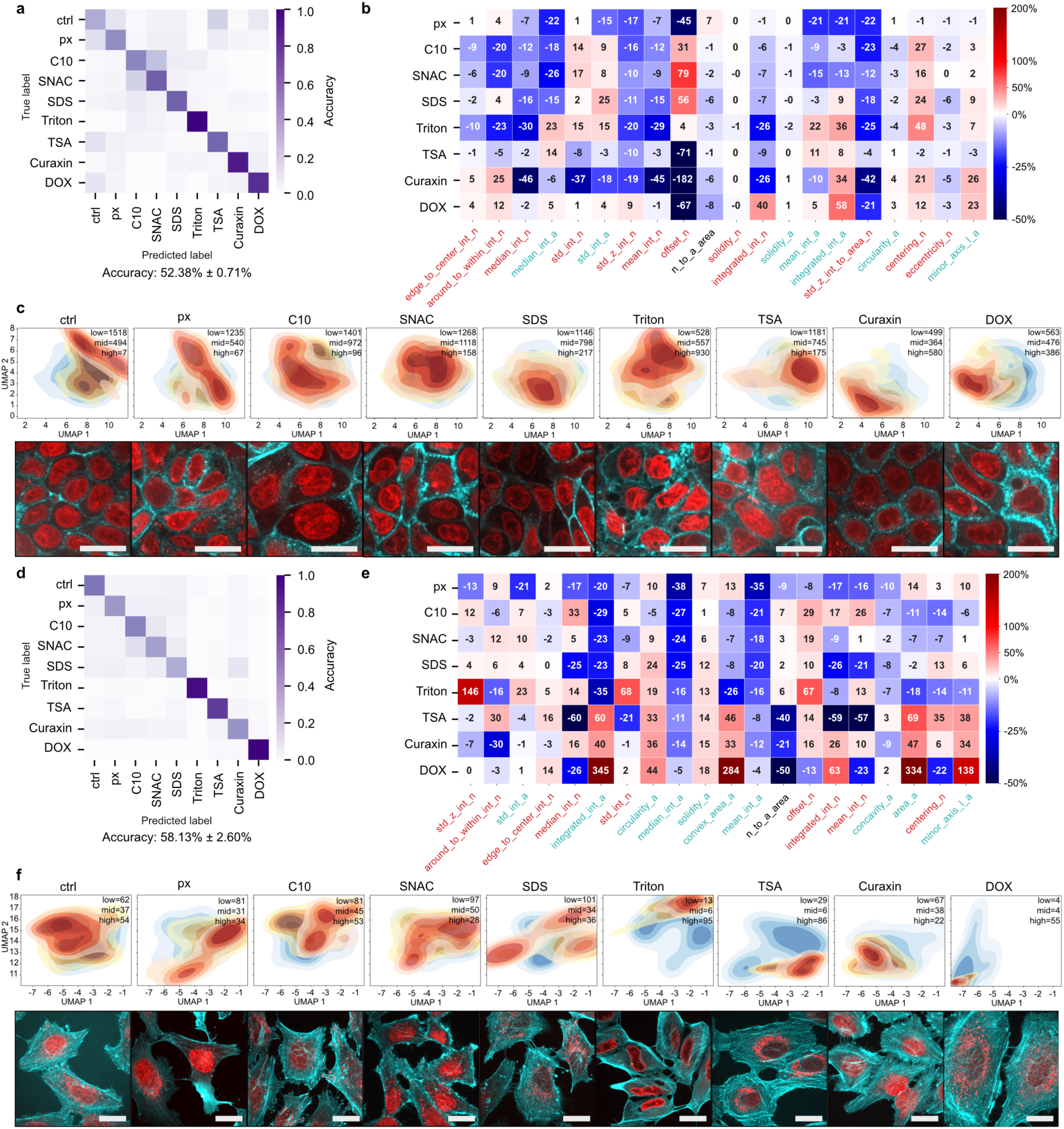
Multiclass classification and cross-cell-type generalization of MORPHIS. (a,d) Confusion matrix of multiclass classification of (a) Caco-2 cells and (d) HeLa cells treated with eight different compounds. Mean accuracy of cross-validation folds is indicated below each matrix. (b,e) Heatmap of percentage change with respect to control (ctrl) of the top 20 SHAP features for (b) Caco-2 and (e) HeLa. Values correspond to feature shifts contributing to class discrimination (c,f) UMAP of feature-space of the extracted features color coded by the probability of the XGBoost classifier for (c) Caco-2 and (f) HeLa cells. The color indicates single-cell probabilities grouped into low response (p < 0.35, blue), mid response (p = 0.35–0.7, yellow), and high response (p ≥ 0.7, red). Underneath is a representative average intensity projection (AIP) of z-stacks (0.5 μm between stacks) of non-fixed Caco-2 cell monolayers, with nucleus in red and actin in cyan for ctrl, 30 min exposure with L-penetramax (px), sodium caprate (C10), sodium N-[8-(2-hydroxybenzoyl) amino] caprylate (SNAC), sodium dodecyl sulfate (SDS), Triton™ X-100 (Triton) and 48h exposure with trichostatin A (TSA), curaxin CBL0137 (Curaxin) and doxorubicin (DOX), showing the unique morphological signature of the condition. (scalebar = 25 µm, Caco-2: i = 4, n = 2, N = 3, HeLa: i = 4, n = 2, N = 2)

**Figure 6:**
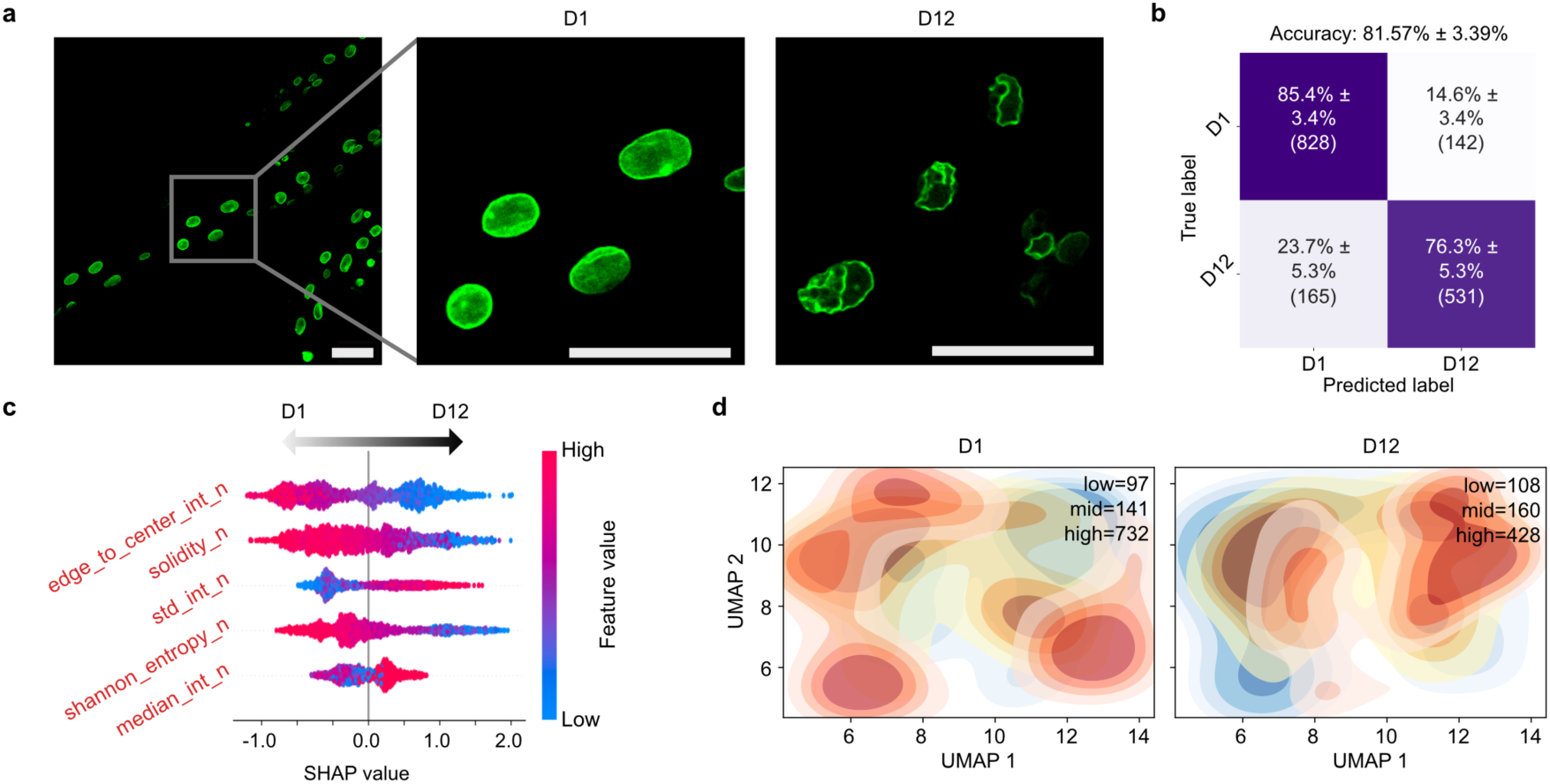
MORPHIS depicts aging-associated nuclear changes of *C. elegans*. (a) Confocal microscopy images of day 1 (D1, young) and day 12 (D12, aged) *C. elegans* nuclei expressing the nuclear lamina reporter LMN-1::GFP. (b) Confusion matrix of the binary classification of young vs. aged population. (c) Beeswarm plot of the top five SHAP-ranked features. (d) UMAP of feature-space of the extracted nucleus features color coded by the probability of the XGBoost classification. The color indicates single-cell probabilities grouped into low response (p < 0.35, blue), mid response (p = 0.35–0.7, yellow), and high response (p ≥ 0.7, red). (scalebar = 25 µm, i = 1, n = 1, N >45)

We first examined treatment-specific morphological responses of Caco-2 monolayer cells under four conditions, one control (ctrl, Fig. 3a) and three compounds representing diverse mechanisms: a senescence-inducing anticancer drug, doxorubicin, producing characteristic morphological damage-induced senescence and nuclear changes (DOX, Fig. 3b), a peptide permeation enhancer L-penetramax (px) (Fig. 3c), and a surfactant Triton™ X-100 (Triton, Fig. 3d), known to adhere to and solubilize the cell membrane, respectively. These perturbations span distinct mechanisms enabling a detailed exploration of the treatment-specific morphological signatures (analyses of all eight treatments can be found in Supplementary Information Fig. 7-21). Prior to analysis, the input segmentation masks were modified based on predefined exclusion criteria to exclude artefacts (Supplementary Information Fig. 9-12).

The pairwise classification of the control and the 48h DOX-treated Caco-2 monolayer cells achieved an accuracy of 80.02±1.07% (Fig. 3e, Supplementary Information Fig. 13a), supporting that MORPHIS can reliably distinguish between the control and the treated population. The top ranked features included nuclear edge-to-center intensity ratio, shannon entropy of the actin and standard deviation of nuclear z-stack intensity (reflecting nucleus elongation/height(Mortensen et al. 2025)) (Fig. 3f). As expected, the features were not affected by technical and biological replicates, signifying their importance in identifying key biological changes (Supplementary Information Fig. 14). A convex hull plot of the top three features shows a clear separation between the classes, illustrating that the top-ranked features not only drive classification but also provide an interpretable representation of the morphological signature (Fig. 3g). DOX-treated cells show a higher edge-to-center nuclear intensity ratio, with more signal at the nuclear periphery than in the center, indicating enhanced peripheral nuclear enrichment. An elevated shannon entropy of the actin reflects greater cytoskeletal heterogeneity while slightly elevated standard deviation of nuclear z-stack intensity is indicative of a more elongated nucleus signal (Fig. 3h). Overall, DOX promotes peripheral nuclear enrichment, nuclear enlargement, and increased cytoskeletal variability consistent with a senescence-associated morphological signature arising from DNA damage-induced stress(Hernandez-Segura et al. 2018; Luzhin et al. 2023; Hu and Zhang 2019). MORPHIS successfully translates complex single-cell morphological responses into quantitative, interpretable signatures.

The permeation enhancer px is believed to interact transiently with the cell membrane(Panou et al. 2023; Diedrichsen et al. 2025), yet MORPHIS reaches an accuracy of 70.55±1.32% (Fig. 3i, Supplementary Information Fig. 13b) for the pairwise comparison of control and px-treated Caco-2 monolayer cells (Fig. 3j). The three top ranked features were standard deviation of nuclear z-intensity, median of the actin intensity, and standard deviation of the nuclear intensity (Fig. 3k). The distributions of the top features (Fig. 3l) show a greater overlap between control and px-treated cells, than for DOX-treated cells, consistent with the lower overall accuracy. In short, px exposure leads to slight nuclear flattening(Panou et al. 2023), without major changes to chromatin organization likely due to loss of epithelium tightness.

Finally, applying MORPHIS to the pairwise classification of control and Triton-treated Caco-2 monolayer cells, resulted in an overall accuracy of 95.69±0.30% (Fig. 3m, Supplementary Information Fig. 13c) highlighting extensive morphological cell perturbation. Triton is extensively used in molecular biology for cell lysis, membrane permeabilization, and is a go-to reagent in protocols involving Western blotting, immunostaining and ELISA, or for inducing apoptosis(Mortensen et al. 2025; Picache et al. 2004). The top features for Triton-treated cells were exclusively nuclear intensity descriptors (Fig. 3n,o) showing centralization of the nuclear signal and increased nuclear heterogeneity, with reduced peripheral enrichment compared to the control (Fig. 3p). These are consistent with pronounced nuclear remodeling associated with apoptotic or cytotoxic responses(Mandelkow et al. 2017; Kang et al. 2025).

By analyzing three treatments with mechanistically distinct perturbations: DNA intercalation (DOX), membrane interaction (px), and permeabilization-induced cytotoxicity (Triton), MORPHIS reveals distinct morphological signatures for each, with top-ranked features corresponding to the biologically relevant cellular responses. This demonstrates that the method captures biologically meaningful, interpretable changes across thousands of single cells independent of origin of morphological change.

### Treatment-dependent heterogeneity in cellular morphological responses

Cell-to-cell heterogeneity is a hallmark of biological responses. Even under identical treatment conditions, cells exhibit variable degrees of morphological change(Niepel et al. 2009) independent of cell density and spatial location (Supplementary Information Fig. 15), typically assessed qualitatively. To quantitatively characterize this variability, we applied MORPHIS to the representative binary classification of control and DOX-treated Caco-2 monolayer cells. UMAP embeddings of the raw single-cell morphological feature-space reveal a gradual shift between the confidently predicted control and DOX-treated cells (Fig. 4a), indicative of a continuum of morphological responses from high to low rather than a strict binary separation. Representative cells spanning high and low response probabilities illustrate this gradient; A high-response DOX-treated cell (p = 0.96, Fig. 4a, top right) exhibits pronounced structural alterations, whereas a low-response DOX-treated cell (p = 0.04, Fig. 4a, bottom right) closely resembles the control phenotype. Within the control population, MORPHIS similarly identifies highly representative control cells (p = 0.93, top left) as well as cells with atypical morphology (p = 0.18, bottom left), whose morphology partially overlaps with the treated cells, illustrating the intrinsic heterogeneity of single-cell morphology. Examination of the numeric top features for these representative cells confirms that low-response DOX-treated cells share nearly identical feature profiles with control cells, while high-response DOX-treated cells resemble atypical control cells located near the phenotypic boundary. These apparent misclassifications originate from the continuum of cellular responses rather than from the presence of discrete classification errors, e.g., a non-responding DOX-treated cell is predicted as control because its morphology resembles the untreated phenotype. MORPHIS not only reveals the existence of this heterogeneity but also allows us to interrogate its underlying causes, linking fractional morphological responses to biologically meaningful features and treatment-specific mechanisms across the population.

To quantify this heterogeneity, cells are separated based on predicted probability. We considered cells with prediction probabilities p ≥ 0.7 as the clearest morphology for each condition, appearing most representative of either the control or the treated phenotype. The continuum observed in the UMAP representation of single-cell prediction probabilities supports this stratification: rather than forming discrete clusters, cells are distributed along a gradient from control-like to strongly treated-like morphology, reflecting the gradual variation in response magnitude across the population. For DOX treatment close to 53.5% of the cell population show p ≥ 0.7 (Fig. 4b), while for px it is only 38.2% (Fig. 4c, Supplementary Information Fig. 16a) consistent with px inducing a more conservative morphological change than DOX as observed in Fig. 3. For Triton exposure, 93.8% of the cell population show p ≥ 0.7 (Fig. 4d, Supplementary Information Fig. 6b) indicative of a pronounced and more uniform morphological shift, consistent with its membrane permeabilizing action. In fact, across all eight treatments, the fraction of cells exhibiting a strong response of p ≥ 0.7 varied considerably depending on the type of exposure reflecting both the differential potency of the treatments and the intrinsic heterogeneity of cellular responses (Fig. 4e, Supplementary Information Fig. 17-21).

Together, these analyses demonstrate that MORPHIS in addition to resolving treatment-specific morphological signatures, provides a quantitative metric of the distribution and magnitude of responses within cell populations. By allowing differentiation of treatments with similar average effects but distinct single-cell responses and by integrating the probability-based classification with feature-level interpretation and low-dimensional embedding, MORPHIS enables quantitative characterization of phenotypic heterogeneity at single-cell resolution.

### MORPHIS supports multiclass classification of both treatment-induced and aging-associated morphological changes *in vitro* and *in vivo*

MORPHIS was applied to a multiclass classification task encompassing all experimental conditions, one control and eight distinct treatments. The confusion matrix for the Caco-2 cells is dominated by diagonal entries, indicating that correct predictions substantially outweigh misclassifications resulting in an overall multiclass accuracy of 52.38±0.71%, close to five times better than a baseline random prediction (Fig. 5a, Supplementary Information Fig. 22). Careful inspection of the confusion matrix also reveals structured patterns of misclassification, highlighting similarities and differences among treatment-induced phenotypes. While the similarity between control and px was evident already in the previous binary classification, the multiclass analysis additionally uncovers a strong resemblance between treatments with either of the two lipidic PEs, sodium caprate (C10) and sodium N-[8-(2-hydroxybenzoyl) amino] caprylate (SNAC). This correspondence aligns with the, to some extent, shared mechanism of action of these compounds in relation to their interaction with the cell membrane leading to increased membrane permeability, supporting the biological relevance of the observed morphological similarities(Twarog et al. 2020). While these similarities result in misclassifications in the confusion matrix (Fig. 5a), they highlight that MORPHIS can pick up similar biologically interpretable morphological effects across the full treatment space.

To gain insight into the morphological characteristics underlying the multiclass predictions, we examined the 20 top-ranked SHAP features (Fig. 5b) and quantified their median percentage changes relative to the control. In Caco-2 cells, 60% of these top-ranked features are associated with nuclear morphology descriptors, predominantly intensity-based measurements. This indicates that treatment-induced morphological changes extend beyond shape alterations to variations in nuclear intensity distributions. The feature heatmap reveals relationships that are similar among compound classes, consistent with the structured patterns in the confusion matrix. Surfactant-interacting lipidic compounds (C10 and SNAC) as well as the non-ionic surfactant (Triton) exhibit broadly consistent feature trends, reflecting their shared ability to perturb cell membranes, whereas the anionic surfactant sodium dodecyl sulfate (SDS) displays a distinct morphological profile likely reflecting a different mode of membrane perturbation involving both lipids and membrane-associated proteins. In contrast, the anticancer drugs (trichostatin A (TSA), curaxin CBL0137 (Curaxin) and DOX) frequently show feature patterns that are anticorrelated with those of surfactants, consistent with their completely different mode of action, that targets intracellular processes (histone deacetylase inhibition, chromatin destabilization and DNA intercalation, respectively) rather than membrane integrity.

Having established population-level trends of the treatment-induced morphological changes, we examined the heterogeneity within individual treatments based on the UMAP embeddings of the multiclass feature-space (Fig. 5c, top). As expected, high-probability (p ≥ 0.7, red) clusters from treatments with similar morphological effects tend to partially overlap. In contrast, cells with morphologically distinct treatments form separate, non-overlapping clusters. For each treatment, cells with the highest predicted single-cell probabilities were used to evaluate representative examples of treatment-specific morphological signatures (Fig. 5c, bottom) linking classification performance to interpretable phenotypic signatures.

Cells with distinct morphologies provide an opportunity to assess whether phenotypes are conserved or cell-type specific and whether they can be accurately resolved by MORPHIS. To address this, we extended the analysis to HeLa cells, which differ fundamentally from Caco-2 cells, particularly when the latter form confluent monolayers. Despite the smaller dataset size, overall classification accuracy was higher for HeLa cells (58.13±2.60%) than for Caco-2 cells, likely reflecting more pronounced and more uniformly treatment-induced morphological changes when cells are not in a constrained monolayer. While the similarity between C10 and SNAC effects is preserved in HeLa cells, the previously observed similarity between control and px is less pronounced (Fig. 5d, Supplementary Information Fig. 23) indicating that certain phenotypic relationships are cell-type dependent.

Inspection of the feature heatmap reveals a shift in dominant descriptors between the two cell types supporting that treatment-induced changes are cell-type specific (Fig. 5e). In HeLa cells actin-related features and global cell shape metrics contribute more prominently to classification compared to Caco-2 cells, where nuclear intensity features dominate. For example, the nuclear height (std_z_int_n) decreases by more than 10% for all treatments except for DOX in Caco-2 cells. In contrast, in HeLa cells, this feature decreases primarily following px exposure but increases markedly (146%) following Triton treatment. Overall, the percentage change observed is more extreme for HeLa cells compared to Caco-2 cells in alignment with the overall higher accuracy displaying a more pronounced morphological change.

Consistent with these observations, UMAP representations show much clearer separation for HeLa cells both for the high (p ≥ 0.7, red) and mid prediction (p = 0.35–0.7, yellow) clusters (Fig. 5f) for several treatments, and in particular for Triton and DOX. By contrast, in Caco-2 cells, cluster separation is less pronounced for these treatments, likely reflecting the more constrained morphology of the monolayer cells, which may limit the magnitude of observable morphological changes. In summary, multiclass classification, feature analysis, and single-cell visualization collectively reveal both conserved and cell-type-specific treatment-induced morphological effects. While there are some similarities in regard to cellular effects e.g. after C10 and SNAC exposure, that are preserved across Caco-2 cells and HeLa cells, other responses, including those to px, differ between the cell types. Feature-level analysis indicates that dominant morphological changes are largely cell-type dependent, with nuclear intensity and cell shape contributing variably across the two cell lines.

To assess MORPHIS on an *in vivo* setting and in a system that is driven by intrinsic biological variation rather than ligand-induced external perturbation, we utilized the well-established aging model in *C. elegans*. MORPHIS was evaluated on a dataset of young (D1) and aged (D12) nuclei of *C. elegans*, expressing a fluorescent reporter of the nuclear lamina protein lamin (LMN-1::GFP) *in vivo* (Fig. 6a, Supplementary Information Fig. 24-25). MORPHIS accurately predicts (81.57±3.39%) the annotated population (Fig. 6b) and the extracted key features captured age-associated changes in nuclear morphology(Pérez-Venteo et al. 2026). Specifically, aged nuclei exhibited a lower nuclear edge-to-center intensity ratio and a lower solidity (Fig. 6c), characterized by nuclear envelope folding and lobulation (Fig. 6a, Supplementary Information Fig. 24). In addition, a higher standard deviation of nuclear intensity arises among aged nuclei, reflecting uneven distribution of nuclear lamina (Fig. 6c and Supplementary Information Fig. 26). This finding is consistent with previous observations that *C. elegans* nuclei undergo age-dependent deterioration with a strong stochastic component(Herndon et al. 2002; Haithcock et al. 2005). UMAP representations of the feature-space (Fig. 6d, Supplementary Fig. 26) further highlight the heterogeneity within the aged population consistent with the diversity in nuclear morphology among aged cells.

Taken together, all these results demonstrate the capacity of MORPHIS to capture both broad trends and, cell-type specific phenotypic responses, highlighting its utility for dissecting complex morphological effects across diverse cellular contexts, while retaining biological interpretability.

## Discussion

Quantitative analysis of cell morphology is central to understanding cellular state, function, and response to various perturbations. Despite great advances in imaging technologies and computational tools, contemporary methods remain constrained by several limitations. Classical feature-based approaches rely on limited geometric features such as area or aspect ratio, reducing complex cellular structure to a small number of descriptors. While intuitive and computationally efficient, these descriptors oversimplify cellular structure and often fail to capture the full spectrum of morphological variation and phenotypic signatures. Deep learning-based models for cell morphology analysis, learn rich and high-dimensional representations directly from images without the need for human predefined features. Convolutional neural networks and related architectures have demonstrated impressive performance in cell classification and while these approaches offer incredible consistency and performance, the representational power of deep learning models comes at the cost of limited interpretability(Wysocka et al. 2023; Wagle et al. 2024; Gao et al. 2025). The latent features learned by these models are typically abstract and not always biology specific. These limitations highlight the need for a quantitative cell morphology framework that balances representational richness with biological interpretability.

MORPHIS addresses this gap by introducing an interpretable feature representation that includes a spectrum of complementary aspects of cellular organization, including shape, intensity distribution, texture, and compartmental architecture. In doing so, it extends above and beyond simple average changes in individual features across the population to an optimized framework that encodes biologically meaningful representation of morphological variation while preserving direct biological traceability of each feature. Importantly, the predictive performance is not achieved at the expense of accuracy. Across multiple classification tasks, MORPHIS performs on par with the state-of-the-art deep learning model scDINO, despite relying on explicitly defined descriptive features rather than learned latent representations. This demonstrates that high classification accuracy in cell morphology does not intrinsically require opaque models, and that interpretable representations can capture much of the relevant biological information. The value of interpretability is illustrated in DOX-specific response where MORPHIS captures the DOX-specific morphological signature through analytically rich while interpretable features, including nuclear enlargement, peripheral nuclear enrichment, and increased cytoskeletal variability associated with DNA damage-induced, senescence-associated structural changes. Rather than providing only a classification outcome, MORPHIS is a highly interpretable platform that links predictive performance to explicit biologically significant phenotypic signature alterations.

Beyond population-level comparisons, MORPHIS enables systematic exploration of single-cell morphological heterogeneity by capturing heterogeneous fractional responses that are typically overlooked by conventional approaches averaging responses across cells. Inherently, some cells display strong alignment with the treated condition, while a fraction of cells may show subtle morphological changes, leading to continuous heterogeneous population-level phenotypes(Snijder et al. 2009; Bray et al. 2016). Because predictions are generated at the level of individual cells using explicit morphological features, the resulting classification prediction probability can be interpreted as a quantitative measure of how closely a given cell resembles its annotated condition. This provides a continuous readout of cellular response, rather than a binary assignment, allowing cells within the same population to be ordered by the extent of their morphological change. This single-cell resolution reveals heterogeneous responses that are distinct to the type of perturbations and while population-level comparisons capture average treatment effects, the distribution of single-cell prediction scores uncovers variability in cellular response magnitude. As a result, our approach supports simultaneous analysis of morphology both at the population-level and single-cell level linking global phenotypic signatures to cell-to-cell variation in morphological response enabling exploration of the differences in key features for cells within and across treatments quantitatively and robustly.

Consistent with its ability to resolve single-cell variability, MORPHIS generalizes across multiple diverse treatments and diverse cell types. The explicit nature of the features allows morphological differences associated with specific treatments to be directly linked to distinct and, importantly, quantifiable aspects of cell morphology for diverse data sets. For example, MORPHIS identified treatment-specific emerging signatures for treated HeLa cells that were absent in the control population. In contrast, treatment of Caco-2 cells did not generate entirely new morphological signatures, rather it shifted the distribution of fractional responses, highlighting cells that exhibited a stronger resemblance to the treated phenotype.

MORPHIS further, effectively captures key nuclear features associated with physiological ageing in an independent dataset derived from young and aged *C. elegans* nematodes demonstrating its applicability beyond externally perturbed systems. This also showcases that MORPHIS can provide a quantitative phenotyping of nuclear aging *in vivo* and its utility to predict and evaluate aging(Papandreou et al. 2023; Wu et al. 2024; López-Otín et al. 2023) and its potential reversal by treatments(Yang et al. 2023).

Together, these findings suggest that MORPHIS enables not only identification of key biological features but also uncovers the fractional response within a treated population. In doing so it offers a practical middle ground between few classical cell shape metrics and deep learning-based approaches by combining generalizability, accuracy, and interpretability in a single framework. Its ability to quantify treatment-induced morphological changes *in vitro* and attain quantitative phenotyping of nuclear aging *in vivo*, demonstrates the robustness and versatility of MORPHIS. Crucially, this enables cell morphology to be used not only as a phenotypic readout at the population level, but also as a source of single-cell mechanistic insight into how perturbations and cellular states manifest in form.

## Methods

### Materials

All chemicals are of analytical grade and purchased from Sigma-Aldrich Denmark (Merck Life Science A/S), unless otherwise stated. Caco-2 cell line (ATCC HTB-37) and HeLa cells line (ATCC (CCL-2)) were purchased from American Type Cell Cultures (ATCC, Manassas, VA, USA). Dulbecco’s Modified Eagles Medium (DMEM), Dulbecco’s Phosphate Buffered Saline (DPBS), trypsin-EDTA, penicillin/streptomycin, L-glutamine, non-essential amino acids (NEAA), and Hank’s Balanced Salt Solution (HBSS) was purchased from Merck Life Science A/S (Søborg, Denmark). Fetal bovine serum (FBS) was acquired from Fisher Scientific (Slangerup, Denmark). Hanks’ Balanced Salt Solution (HBSS), without divalent cations was purchased from Capricorn Scientific GmbH (Hessen, Germany). L-penetramax (px, KWFKIQMQIRRWKNKR, >98% purity) was purchased from Synpeptide (Shanghai, China). Hydroxyethyl piperazineethanesulfonic acid (HEPES), sodium caprate (C10, CAS: 1002-62-6), sodium N-[8-(2-hydroxybenzoyl) amino] caprylate (SNAC, CAS: 203787-91-1), sodium dodecyl sulfate (SDS, CAS: 151-21-3), trichostatin A (TSA, T1952), doxorubicin (DOX, D5220), Triton^TM^ X-100 (Triton) were purchased from Merck Life Science A/S (Søborg, Denmark). Curaxin CBL0137 (Curaxin) was purchased from MedChemExpress (Sollentuna, Sweden). MTS-PMS CellTiter 96® AQueous One Solution Cell Proliferation Assay was acquired from Promega (Madison, WI, USA). NucRed^TM^ Live 647 ReadyProbes^TM^ Reagent and CellMask^TM^ Actin Stains were purchased from Thermo Fisher Scientific (Waltham, MA, USA).

### *In vitro* cell culturing

The Caco-2 human colorectal adenocarcinoma cell line was maintained in T75 cm² flasks (Corning, Merck Life Science A/S) using DMEM supplemented with 10% (v/v) FBS, 2 mM L-glutamine, 90 IU/mL penicillin, 90 μg/mL streptomycin and 0.1 mM NEAA. Cells were cultured in a humidified incubator (5% CO₂, 95% O₂, 37°C). The culture medium was replaced every other day, and cells were passaged weekly using trypsin-EDTA upon reaching 90% confluency.

For viability assessment, Caco-2 cells were seeded at a density of 30,000 cells/well in clear 96-well plates (BRANDplates® cellGrade™, Sigma-Aldrich). The cells were cultured for 6-8 days in complete DMEM, with medium change every other day. Cells from passages 4 to 7 were used for viability experiments.

For microscopy experiments, Caco-2 cells were seeded at a density of 1 × 10⁵ cells/well in a µ-Slide 8 Well (Cat.No: 80826, ibiTreat, Ibidi GmbH, Germany). The cells were cultured for 6-8 days in ibidi-slides (5% CO₂, 95% O₂, 37°C) in complete DMEM, with medium change every other day. Cells from passages 7 to 11 were used for microscopy experiments.

The HeLa cells were maintained in T25 cm² flasks (Corning, Merck Life Science A/S) using DMEM supplemented with 10% (v/v) FBS, 2 mM L-glutamine, 90 IU/mL penicillin, 90 μg/mL streptomycin and 0.1 mM NEAA. Cells were cultured in a humidified incubator (5% CO₂, 95% O₂, 37°C). The culture medium was replaced every other day, and cells were passaged using trypsin-EDTA before reaching 90% confluency twice a week.

For viability assessment, HeLa cells were seeded at a density of 3,000 cells/well in clear 96-well plates (BRANDplates® cellGrade™, Sigma-Aldrich). The cells were cultured for 3 days in complete DMEM, with medium change every other day. Cells from passages 13-15 were used for the viability experiments.

For microscopy experiments, HeLa cells were seeded at a density of 5000 cells/well in a µ-Slide 8 Well (Cat.No: 80826, ibiTreat, Ibidi GmbH, Germany). The cells were cultured for 3 days in ibidi-slides (5% CO₂, 95% O₂, 37°C) in complete DMEM, with medium change every other day. Cells from passages 17-19 were used for microscopy experiments.

### Viability assay

The metabolic activity was assessed after 30 min exposure for buffer (10 mM HEPES in HBSS pH 7.4 (hHBSS)), penetramax, C10, SNAC, SDS and Triton and after 24h, 48h and 72h for TSA, Curaxin and DOX. For the 30 min exposure the test preparations were made in hHBSS (hHBSS w/o divalent cations for C10) while for 24h-72h exposure, the test solutions were made in complete DMEM. The test solutions were removed prior to incubation with the MTS-PMS reagent (240 μg/mL MTS and 4.8 μg/mL PMS in hHBSS) for 1-2h before measuring absorbance at 492nm and 690 nm. The normalized viability was calculated as reported in (Mortensen et al. 2025).

### Confocal Spinning Disc Microscopy imaging

Caco-2 monolayers grown for 6-8 days in ibidi-slides were washed with 10 mM HEPES in HBSS pH 7.4 (hHBSS) and stained with a solution of NucRed^TM^ Live 647 ReadyProbes^TM^ Reagent (2 drops pr mL, red 640 nm excitation) and CellMask^TM^ Actin Stains (1×, cyan 488 nm excitation) in hHBSS (pH = 7.4) and incubated for 20 min at 37°C. The staining solution was removed after 20 min, and the cells were washed with hHBSS twice to remove the staining solution. HeLa cellswere grown for 3 days in ibidi-slides before imaging and only incubated with the staining solution for 5 min. 280 µL of fresh hHBSS was added to each chamber and the cells were incubated with test solutions for 30 min before imaging (hHBSS w/o divalent cations for C10). For TSA, Curaxin and DOX, the 48h incubation in complete DMEM took place before staining. Before imaging, the test solutions were removed and cells were washed twice with hHBSS.

Imaging was performed with an inverted spinning disk confocal microscope (Olympus SpinSR10, Olympus, Tokyo, Japan) equipped with a 60× oil immersion objective with a numerical aperture of 1.42 (UPLXAPO60XO, Olympus, Evident, Tokyo, Japan) and CMOS camera (PRIME 95B, Teledyne Photometrics, Tucson, AZ, USA).

Z-stacks of 151 stacks with a step size of 0.5 µm for each frame was acquired with 20% laser power for 488 laser excitation and 20% laser power for 640 nm laser excitation with 100 ms exposure time at 37°C. For each condition at least 4 videos at different positions in each well were collected (i > 4, n = 2, N = 2-4). Here, i represents the number of videos, n denotes the technical replicates and N the biological replicates.

### *C. elegans* maintenance and imaging

The lamin reporter strain LW697 (*ccIs4810 [(pJKL380.4) lmn-1p::lmn-1::GFP::lmn-1 3’utr + (pMH86) dpy-20(+)] I*) was obtained from the *Caenorhabditis Genetics Center.* The animals were maintained in nematode growth media plates seeded with OP50 bacteria at 20°C. Imaging was carried out using hermaphrodite worms. At the indicated days of adulthood, worms were mounted in 2% agarose pads with 20mM tetramisole in M9 buffer. Image acquisition was performed at the hypodermal plane of the posterior part (between vulva and tail) of the animals using a ZEISS LSM900 Axio Observer.Z1/7 confocal microscope equipped with an LD LCI Plan-Apochromat 40x/1.2 Imm Korr DIC M27 objective and an Airyscan 2 detector.

### Image analysis

Quantitative image analysis was performed using in-house software in Python 3.10 and outlined in the following paragraphs.

### Segmentation

The raw z-stack images were pre-processed before segmentation. We automatically extracted the average 640 nm intensity in each z-plane and found the maximum value representing the center of the nuclei layer. We then constructed the average intensity projection (AIP) of both channels of the median z-plane ± 3 z-planes resulting in an AIP of 7 z-planes in the center of the nucleus layer. An AIP was chosen to reduce noise, while maintaining a relatively thin slice of the cell layer. We also excluded pixels with an intensity of 400 (A.U.) higher than the mean intensity of the full image for the segmentation, to exclude bias from high intensity artifacts.

All AIP images were segmented in Cellpose 3.0 using the Cyto model for the actin channel and nucleus model for the nucleus channel, individually. All images were manually evaluated, and changes were made to the segmented mask if necessary (Supplementary Information Fig. 9).

For *C. elegans,* segmentation was performed on a single-slice image using the nucleus model for the nucleus channel and evaluated manually after segmentation.

### MORPHIS

MORPHIS is a machine learning framework developed to extract mechanisms of cellular morphological perturbations. It consists of two modules; one for extraction of single-cell features to define the morphological signature and one for analyzing the feature distributions by classification, ranking and subsequent statistical analysis.

### Feature extraction

Prior to feature extraction, the output masks for the 640 nm channel and 488 nm channel from Cellpose were paired for the same field-of-view. To ensure that a dataset consisted of complete dual-segmented cells for analysis, any unpaired nucleus and actin masks were removed with the requirement, that at least 90% of the nucleus mask must be contained within the coordinates of the actin mask to make a pair. Cells with only partial or faulty segmentation were excluded from analysis and for Caco-2 images, segmentation masks were removed if they had one or more pixels on the edge of the field of view (Supplementary Information Fig. 10-12).

Lastly, the z-stack images were normalized to the 99^th^ percentile to reduce the effect of bright artifacts for a non-biased comparison of intensity-based features across different images.

Features were extracted for each individual mask pair using all points on the boundary and the interior of the mask shapes in combination with the normalized pixel intensity values (See Supplementary Information Table 1 for the full list of features and descriptions). For the shannon entropy the pixels within a mask were denoised with a Gaussian filter with a sigma = 1, to mitigate the effect of noise on the feature. Features related to a height estimate of the nucleus were calculated by fitting a gaussian to the mean intensity of pixels within the nucleus mask and extracting the standard deviation. We evaluated the reproducibility of the extracted features across technical and biological replicates (Supplementary Information Fig. 14), and found little variation.

For *C. elegans,* actin-related features and features related to the z-stack were omitted.

### Feature Preprocessing

Prior to model training, all features were standardized to zero mean and unit variance using statistics computed exclusively from the training data within each cross-validation fold. This prevented information leakage between training and validation sets and enabled fair comparison across models with different inductive biases. No additional feature selection or dimensionality reduction was applied unless explicitly stated.

### Model Training, Selection, and Evaluation

Model performance was evaluated on binary and multiclass classification tasks using five-fold cross-validation. For each fold, models were trained on 80% of the data and evaluated on the remaining 20%, with classification accuracy used as the primary performance metric. Hyperparameters for each classifier were optimized independently prior to evaluation using Bayesian optimization implemented with Optuna. For each model, 100 optimization trials were conducted, with search spaces manually defined based on model-specific parameters while the model that was used for the feature importance calculation was trained on all the data. All models were trained, optimized, and evaluated under identical preprocessing, cross-validation, and optimization protocols to ensure a fair and controlled benchmarking framework.

Among the evaluated classifiers(Support Vector Machine, tree-based and logistic classifier, Supplementary Information Fig. 1), the tree-based XGBoost model consistently demonstrated superior performance across cross-validation folds and was therefore selected as the primary model for downstream analyses and to illustrate the analytical workflow presented in this study. Final model accuracies were computed as the mean classification accuracy across all cross-validation folds and the respective standard deviation of accuracy for error bars.

### UMAP-Based Probability Aggregation

To obtain prediction probabilities for all samples without restricting analysis to a single held-out test set, prediction probabilities were aggregated across cross-validation folds. For each fold, the trained model was used to generate class probabilities for the corresponding validation split. These probabilities were iteratively appended across all folds, yielding prediction probabilities for every data point in the dataset. This strategy enabled downstream visualization and probabilistic analysis, including UMAP embedding of prediction probabilities, while preserving strict separation between training and validation data at every stage.

### Feature Importance and Model Interpretability

To quantify feature contributions and enable model interpretability, feature importance was assessed using SHAP (SHapley Additive exPlanations) values for all evaluated classifiers. SHAP values were computed using explainer methods appropriate to each model class: a TreeExplainer was used for tree-based models, while a KernelExplainer was applied to non-tree-based models using sampled background data and subsampling of explained instances to ensure computational tractability.

To characterize global feature relevance within the context of model learning, SHAP values were computed using a final model trained on the full dataset with the five-fold Optuna optimized hyperparameters. This approach does not affect model selection or performance estimation, which were exclusively based on cross-validation, and was chosen as such to investigate the underlying information in the data that is important for the separation of the given classification classes rather than a test of generalization of the model while enabling us to have a full data distribution. Feature importance rankings were derived from the mean absolute SHAP value across samples, reflecting the overall contribution of each feature to the model’s predictions.

### External benchmarking

Three published methods were used to benchmark our method: VAMPIRE, CAJAL and scDINO. For all of the methods cell masks on the boundaries of the images were excluded so the input data are compatible with the performance of our proposed method and XGBoost was selected, and hyperparameter optimized separately for each one of the methods following identical protocol with our method.

#### VAMPIRE

100 contour points were extracted per valid cell mask was extracted to pass as features to an XGBoost classifier which was optimized using Optuna identically with the classifier of our proposed method. The performance metric is the mean accuracy of five-fold cross-validation on the whole dataset. VAMPIRE method also outputs six morphological features which were used for benchmarking separately to the outputted contours.

#### CAJAL

50 contour points were extracted to create a Nx100 feature dataset where N is the number of valid cells. CW distances were calculated and 20 dimensions of UMAP were selected as a dataset to pass onto the XGBoost model which again was optimized with Optuna and performance benchmarked on the mean accuracy of the five cross-validation sets.

#### scDINO

The original two-channel images were cropped on a per cell basis with windows of size 300×300 pixels, so all cells are entirely contained to the cropped windows. The cropped images were preprocessed by resizing them to the input size 224×224 and soft masked (replace all pixel values outside the binary mask with the median value of the pixels outside of the masks but inside the crops). The Dinov1-S with 26M parameters was used as provided by their Github repository that was trained on Imagenet data and takes arbitrary number of channels, which for our dataset is two (cytosol and nucleus). For each cropped image, the cls token was extracted and it was used to train and hyperparameter optimize an XGBoost classifier. The mean 5 cross validation accuracy is provided similarly to all the other methods.

### Statistics and reproducibility

All values reported are an average of at least two technical replicates for each biological replicate, unless otherwise stated (n = 2, N = 2-4).

## Acknowledgements

The work was financially supported by the Novo Nordisk Foundation via the Grand Challenge Programs in support of the Center for Optimized Oligo Escape and Control of Disease (NNF23OC0081287), the Center for Biopharmaceuticals and Biobarriers in Drug Delivery (BioDelivery, NNF16OC0021948) and the Center for 4D cellular dynamics (NNF22OC0075851). The work was also financially supported by the Swiss National Science Foundation (310030M_204518) and Villum Foundation Synergy grant (40578). G.K. and N.T. were supported through the European Commission Research Executive Agency Excellence Hub grant “CHAngeing” (Grant Agreement no.: 101087071). The funding bodies were not involved in the work.

## Author contributions

N.S.H. designed the study with F.B. and H.M.N.. F.B. performed *in vitro* cell culturing, viability assessment and microscopy experiments with help from J.S.M. and E.M.N. F.B., A.O. and E.M.N. developed the framework. E.M.N. and A.O. carried out data analysis and did the interpretation with F.B.. A.O. did all benchmarking of the method. G.K. performed experiments and microscopy on *C. elegans* under the supervision of N.T.. F.B. wrote the manuscript with input from all coauthors. All authors discussed the data, interpretation, and edited the final manuscript. N.S.H. had the overall management and supervision of the project with inputs from H.M.N..

## Competing interests

N.S.H. is the CSO and co-founder of EDGE Biotechnologies.

## Data availability

The authors declare that the data supporting the findings of this study are available within the paper and its Supplementary Information files. All manuscript data will be available upon publication to University of Copenhagen ERDA repository.

## Code availability

Code for MORPHIS will be fully available on Hatzakislab github after peer review acceptance.

## Notes

### Competing Interest Statement

Nikos S hatzakis is CSO and co-founder of EDGE Biotechnologies

