## supplementary information for "MORPHIS (MORPHological Interpretable Signature) captures heterogeneous treatment- and aging-related responses of single cells"

*These authors contributed equally

| No. | Compartment | Feature | Sub-set | Description |
|---|---|---|---|---|
| 1 | Nucleus | Area | Size | No. of pixels in the region (scaled) |
| 2 | Nucleus | Convex area | Size | Area of the convex hull (scaled) |
| 3 | Nucleus | Perimeter | Size | No. of horizontal and vertical connections between adjacent border pixels (scaled) |
| 4 | Nucleus | Minor axis length | Size | Assuming ellipse, length of minor axis (scaled) |
| 5 | Nucleus | Major axis length | Size | Assuming ellipse, length of major axis (scaled) |
| 6 | Nucleus | Std Z stack intensity | Size | SD of Gaussian fitted to the mean z intensity of the region |
| 7 | Nucleus | Solidity | Shape | Area/area of convex hull |
| 8 | Nucleus | Eccentricity | Shape | Assuming ellipse, focal distance/major axis length |
| 9 | Nucleus | Elongation | Shape | Major axis length/minor axis length |
| 10 | Nucleus | Circularity | Shape | 4*area/perimeter$^2$ |
| 11 | Nucleus | Variance | Shape | Variance of distances from centroid to contour edge |
| 12 | Nucleus | Concavity | Shape | No. of times the curvature of the region changes sign, normalized with no. of boundary points |
| 13 | Nucleus | Protrusions | Shape | No. of local maxima for curvature of region |
| 14 | Nucleus | Mean intensity | Intensity | Mean of pixel values in region |
| 15 | Nucleus | Median intensity | Intensity | Median of pixel values in region |
| 16 | Nucleus | Integrated intensity | Intensity | Sum of pixel values in region |
| 17 | Nucleus | Std intensity | Texture | SD of pixel values in region |
| 18 | Nucleus | Edge/center intensity | Texture | Mean intensity near edge of region/mean intensity of remaining pixels |
| 19 | Nucleus | Around/within intensity | Texture | Mean intensity of pixels within the actin mask and not the nucleus mask divided by the mean intensity of pixels within the nucleus mask |
| 20 | Nucleus | Shannon entropy | Texture | Shannon entropy (S) of region: S = -sum(pk*log(pk)), pk is the probability of pixel value k |
| 21 | Nucleus | Zstack/area | Spatial | SD of Z stack intensity / area of nucleus |
| 22 | Nucleus | Offset from monolayer | Spatial | The absolute difference between the mean z-slice intensity of the ROI and the mean z-slice intensity of the full FOV, multiplied by the spacing between z-scans to obtain units of length |
| 23 | Nucleus & actin | Centering of nucleus | Spatial | Euclidian distance between nucleus centroid and actin centroid |
| 24 | Nucleus & actin | Nucleus/actin area | Spatial | Area of nucleus region / area of actin region |
| 25 | Actin | Area | Size | No. of pixels in the region (scaled) |
| 26 | Actin | Convex area | Size | Area of the convex hull (scaled) |
| 27 | Actin | Perimeter | Size | No. of horizontal and vertical connections between adjacent border pixels (scaled) |
| 28 | Actin | Minor axis length | Size | Assuming ellipse, length of minor axis (scaled) |
| 29 | Actin | Major axis length | Size | Assuming ellipse, length of major axis (scaled) |
| 30 | Actin | Solidity | Shape | Area/area of convex hull |
| 31 | Actin | Eccentricity | Shape | Assuming ellipse, focal distance/major axis length |

| | | | | |
|---|---|---|---|---|
| 32 | Actin | Elongation | Shape | Major axis length/minor axis length |
| 33 | Actin | Circularity | Shape | 4*area/perimeter $^2$ |
| 34 | Actin | Variance | Shape | Variance of distances from centroid to contour edge |
| 35 | Actin | Concavity | Shape | No. of times the curvature of the region changes sign, normalized with no. Of boundary points |
| 36 | Actin | Protrusions | Shape | No. of local maxima for curvature of region |
| 37 | Actin | Mean intensity | Intensity | Mean of pixel values in region |
| 38 | Actin | Median intensity | Intensity | Median of pixel values in region |
| 39 | Actin | Integrated intensity | Intensity | Sum of pixel values in region |
| 40 | Actin | Std intensity | Texture | SD of pixel values in region |
| 41 | Actin | Shannon entropy | Texture | Shannon entropy (S) of region: S = -sum(pk*log(pk)), pk is the probability of pixel value k |

**Table 1:** Description of each morphological signature feature. (scaled) denotes features where area or lengths are scaled by the physical size of the pixel.

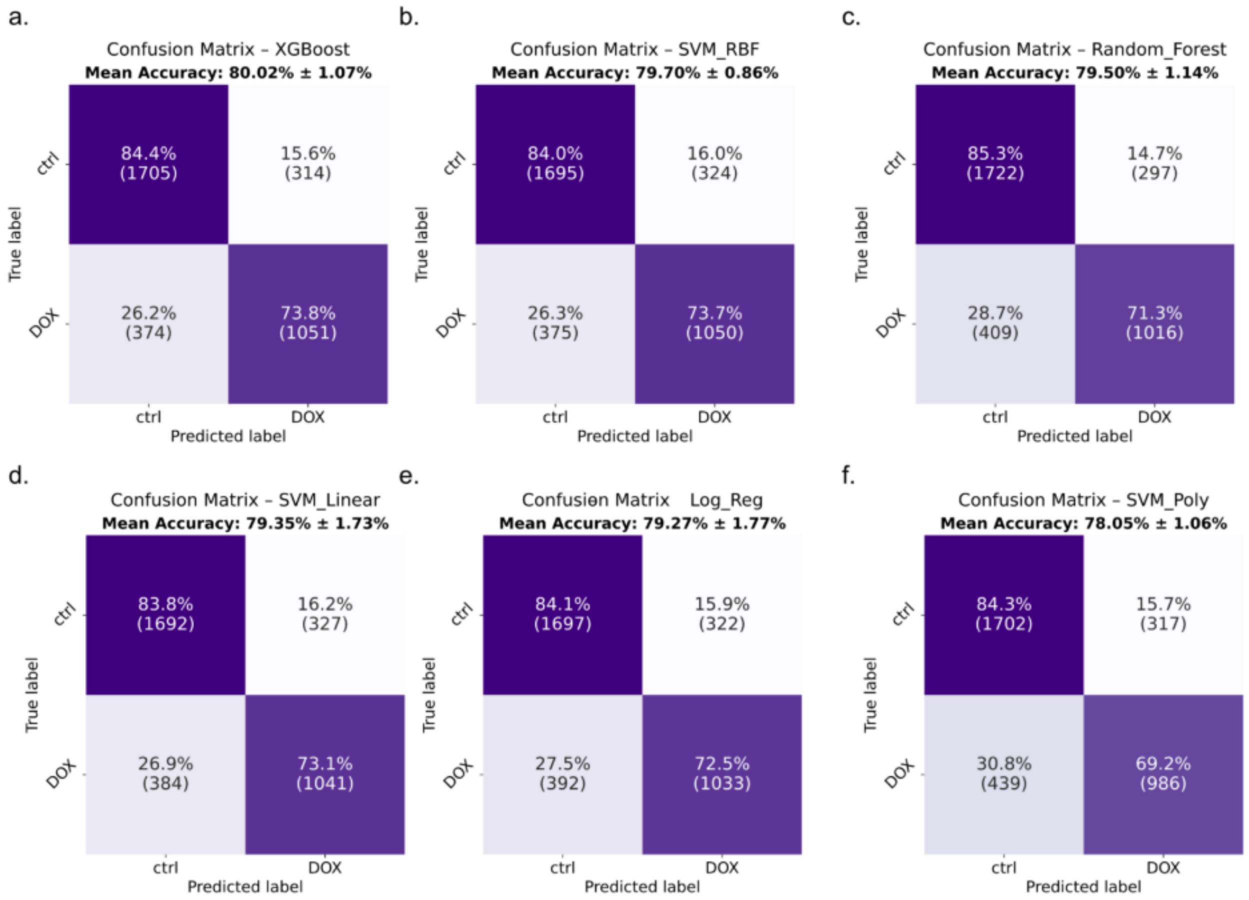


**Figure 1:** Internal validation of MORPHIS. Confusion matrix for the stratified five-fold cross validated classification of control (ctrl) and doxorubicin (DOX) treated Caco-2 monolayer cells for different classifiers: (a) XGBoost, (b) support vector machine with radial basis kernel (SVM_RBF), (c) random forest (Random_Forest) (d) support vector machine with linear kernel (SVM_Linear), (e) logistic regression (Log_Reg) and (f) support vector machine with polynomial kernel (SVM_Poly).

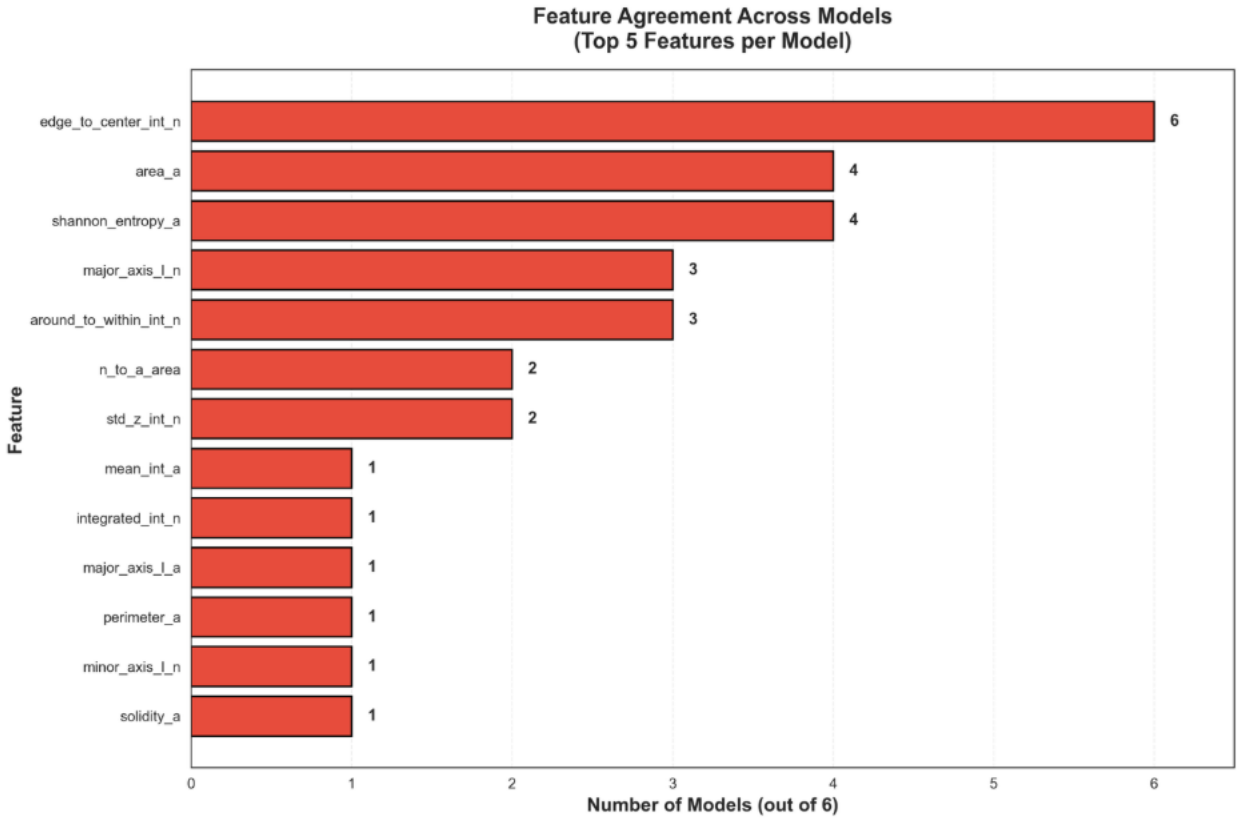


**Figure 2:** Feature agreement across the different models for the binary classification of control (ctrl) and doxorubicin (DOX) treated Caco-2 monolayer cells. Each bar denotes how many models voted for the specific feature in their top 5 features.

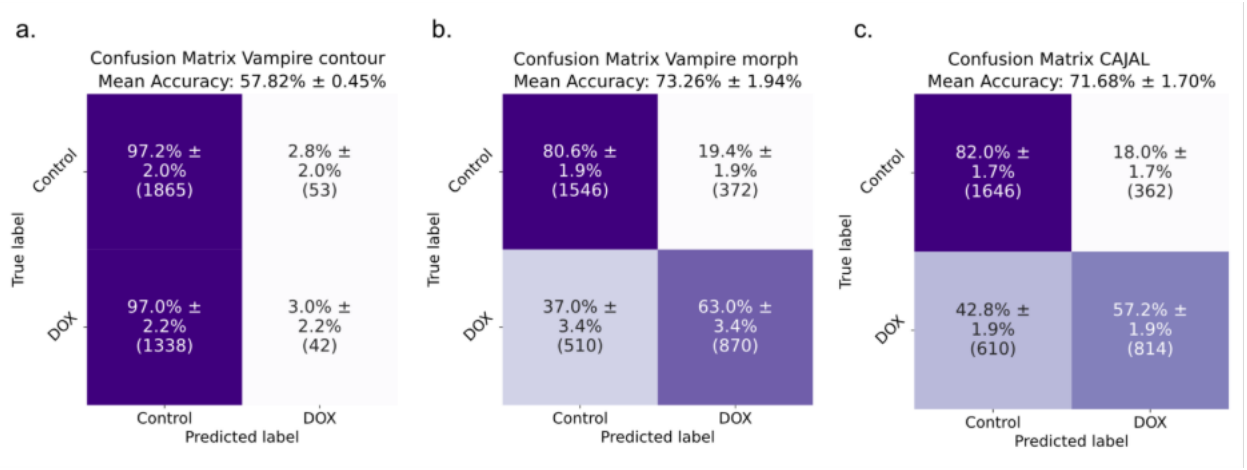


**Figure 3:** Benchmarking against Vampire and CAJAL. Confusion matrix for the stratified five-fold cross validated XGBoost classification of control (ctrl) and doxorubicin (DOX) treated Caco-2 monolayer cells using (a) Vampire contour, (b) Vampire morph and (c) CAJAL to extract features.

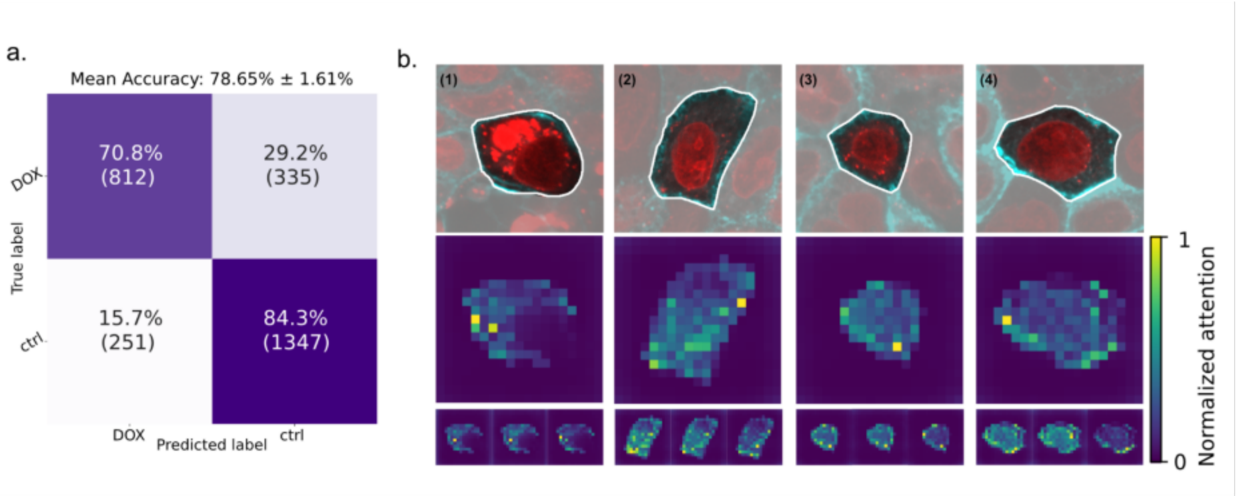


**Figure 4: Benchmarking against scDINO.** (a) Confusion matrix for the stratified five-fold cross validated classification of control (ctrl) and doxorubicin (DOX) treated Caco-2 monolayer cells using scDINO and (b) corresponding attention maps.

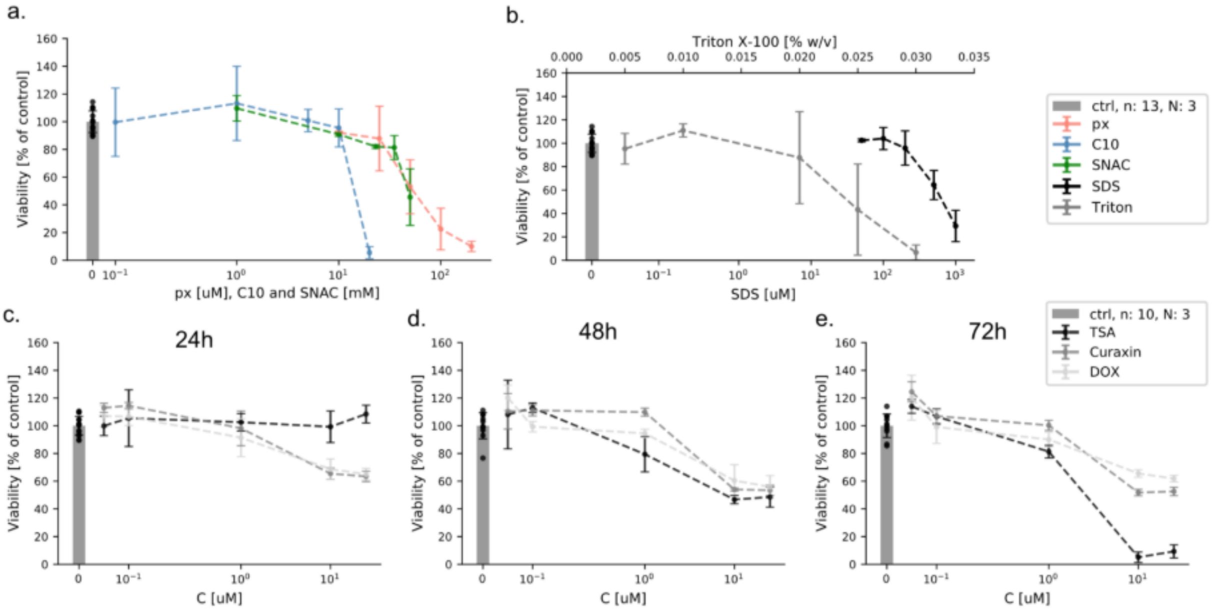


**Figure 5:** MTS-PMS viability [% with respect to control] for Caco-2 cells in 96-well plates for exposure to buffer and different concentration of test-compound. (a) Exposure to hHBSS (ctrl), L-penetramax (px), sodium caprate (C10), sodium N-[8-(2-hydroxybenzoyl) amino] caprylate (SNAC), for 30 min. (b) Exposure to hHBSS (ctrl), sodium dodecyl sulfate (SDS) and Triton™ X-100 (Triton) for 30 min. (c,d,e) Exposure to DMEM+ (ctrl), trichostatin A (TSA), curaxin CBL0137 (Curaxin) and doxorubicin (DOX) for 24h (c), 48h (d) and 72h (e). All points show the mean value for 3 biological replicates. For ctrl the results are from 3 biological replicates and up to 13 technical replicates in total.

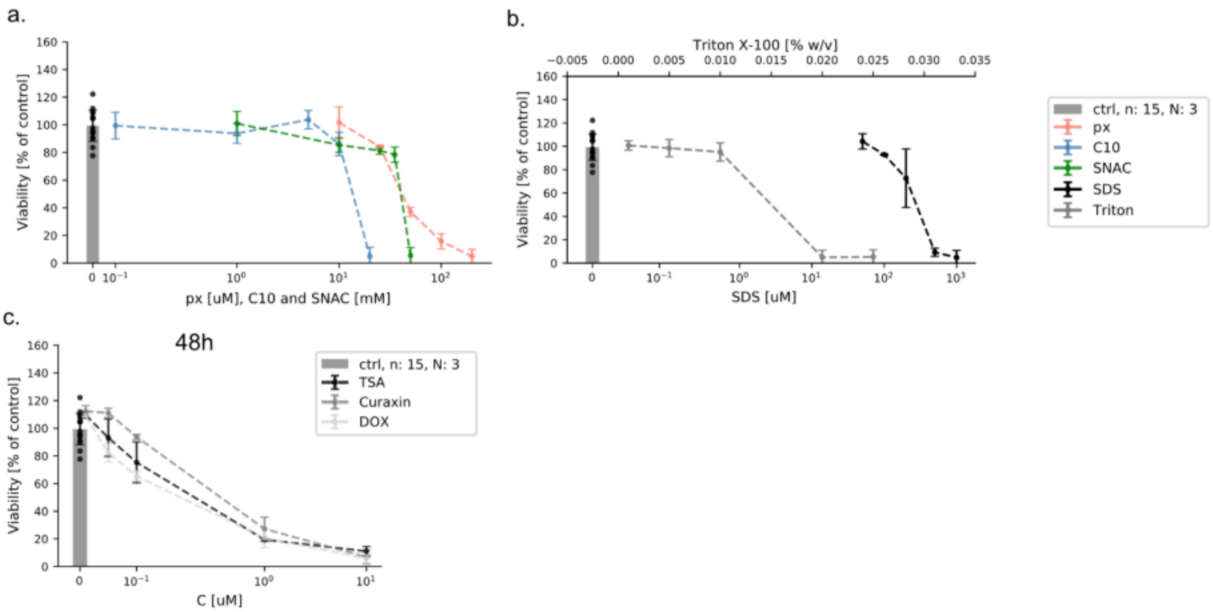


**Figure 6:** MTS-PMS viability [% with respect to control] for HeLa cells in 96-well plates for exposure to buffer and different concentration of test-compound. (a) Exposure to hHBSS (ctrl), L-penetramax (px), sodium caprate (C10), sodium N-[8-(2-hydroxybenzoyl) amino] caprylate (SNAC), for 30 min. (b) Exposure to hHBSS (ctrl), sodium dodecyl sulfate (SDS) and Triton™ X-100 (Triton) for 30 min. (c,d,e) Exposure to DMEM+ (ctrl), trichostatin A (TSA), curaxin CBL0137 (Curaxin) and doxorubicin (DOX) for 48h. All points show the mean value for 3 biological replicates. For ctrl the results are from 3 biological replicates and 15 technical replicates in total.

| | px | C10 | SNAC | SDS | Triton | TSA | Curaxin | DOX |
|---|---|---|---|---|---|---|---|---|
| Caco-2 | 20 µM | 10 mM | 25 mM | 400 µM | 0.02 % (w/v) | 1 µM | 6 µM | 3 µM |
| HeLa | 25 µM | 10 mM | 25 mM | 200 µM | 0.012 % (w/v) | 0.1 µM | 0.2 µM | 0.05 µM |

**Table 2:** Final concentrations used in the microscopy experiments for Caco-2 and HeLa cells, respectively. The concentrations are those that produce 80% viability in the MTS-PMS viability assay after 30 min exposure for permeation enhancer and surfactants: L-penetramax (px), sodium caprate (C10), sodium N-[8-(2-hydroxybenzoyl) amino] caprylate (SNAC), sodium dodecyl sulfate (SDS) and Triton™ X-100 (Triton) and 48h exposure to anticancer drug trichostatin A (TSA), curaxin CBL0137 (Curaxin) and doxorubicin (DOX).

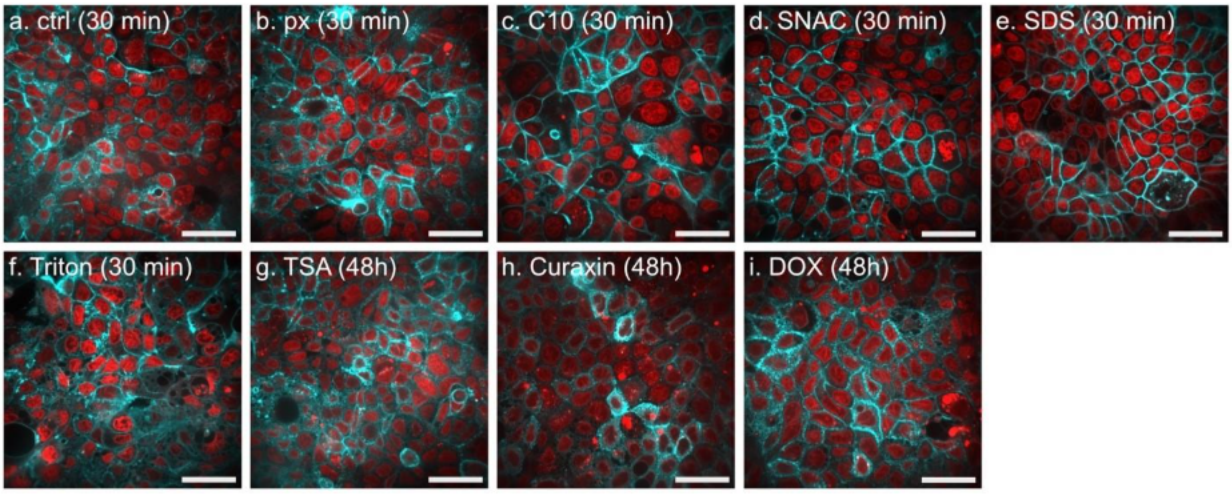


**Figure 7:** Representative average intensity projection (AIP) of z-stacks (0.5 µm between stacks) of live Caco-2 monolayer cells with exposure for 30 min with (a) buffer (hHBSS, ctrl), (b) 20 µM L-penetramax (px), (c) 10 mM sodium caprate (C10), (d) 25 mM sodium N-[8-(2-hydroxybenzoyl) amino] caprylate (SNAC), (e) 400 µM sodium dodecyl sulfate (SDS), (f) 0.02% (w/v) Triton™ X-100 (Triton), and for 48h with (g) 1 µM trichostatin A (TSA), (h) 6 µM curaxin CBL0137 (Curaxin) and (i) 3 uM doxorubicin (DOX). Four field-of-views pr technical replicate were acquired. (cyan: actin, red: nucleus, scalebar = 50 µm, i = 4, n = 2, N = 3).

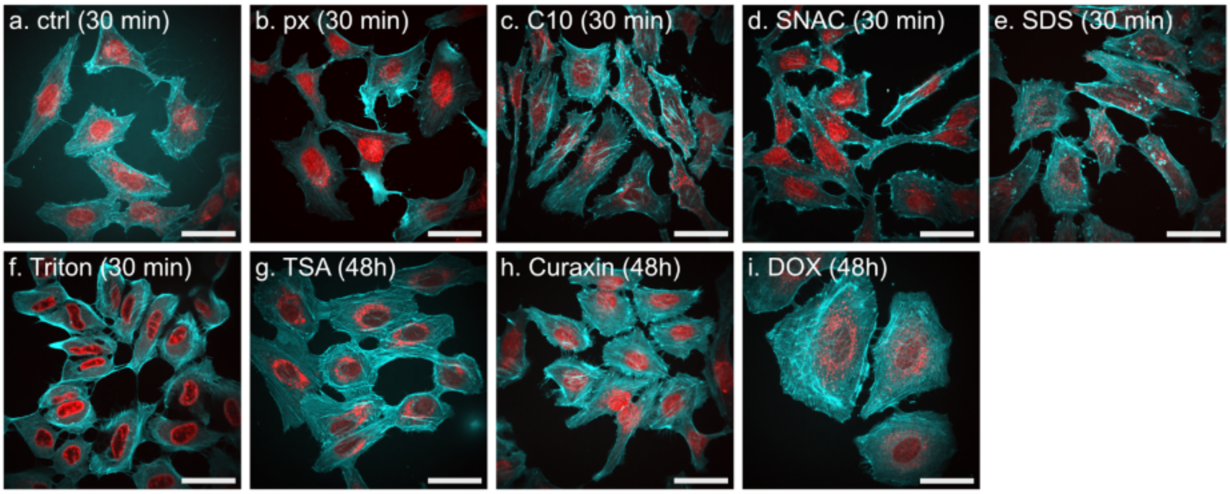


**Figure 8:** Representative average intensity projection (AIP) of z-stacks (0.5 µm between stacks) of live HeLa cells with exposure for 30 min with (a) buffer (hHBSS, ctrl), (b) 25 µM L-penetramax, (px) (c) 10 mM sodium caprate (C10), (d) 25 mM sodium N-[8-(2-hydroxybenzoyl) amino] caprylate (SNAC), (e) 200 µM sodium dodecyl sulfate (SDS), (f) 0.012% (w/v) Triton™ X-100 (Triton) and for 48h with (g) 0.1 µM trichostatin A (TSA), (h) 0.2 µM curaxin CBL0137 (Curaxin) and (i) 0.05 uM doxorubicin (DOX). Four field-of-views pr technical replicate were acquired. (cyan: actin, red: nucleus, scalebar = 50 µm, i = 4, n = 2, N = 2).

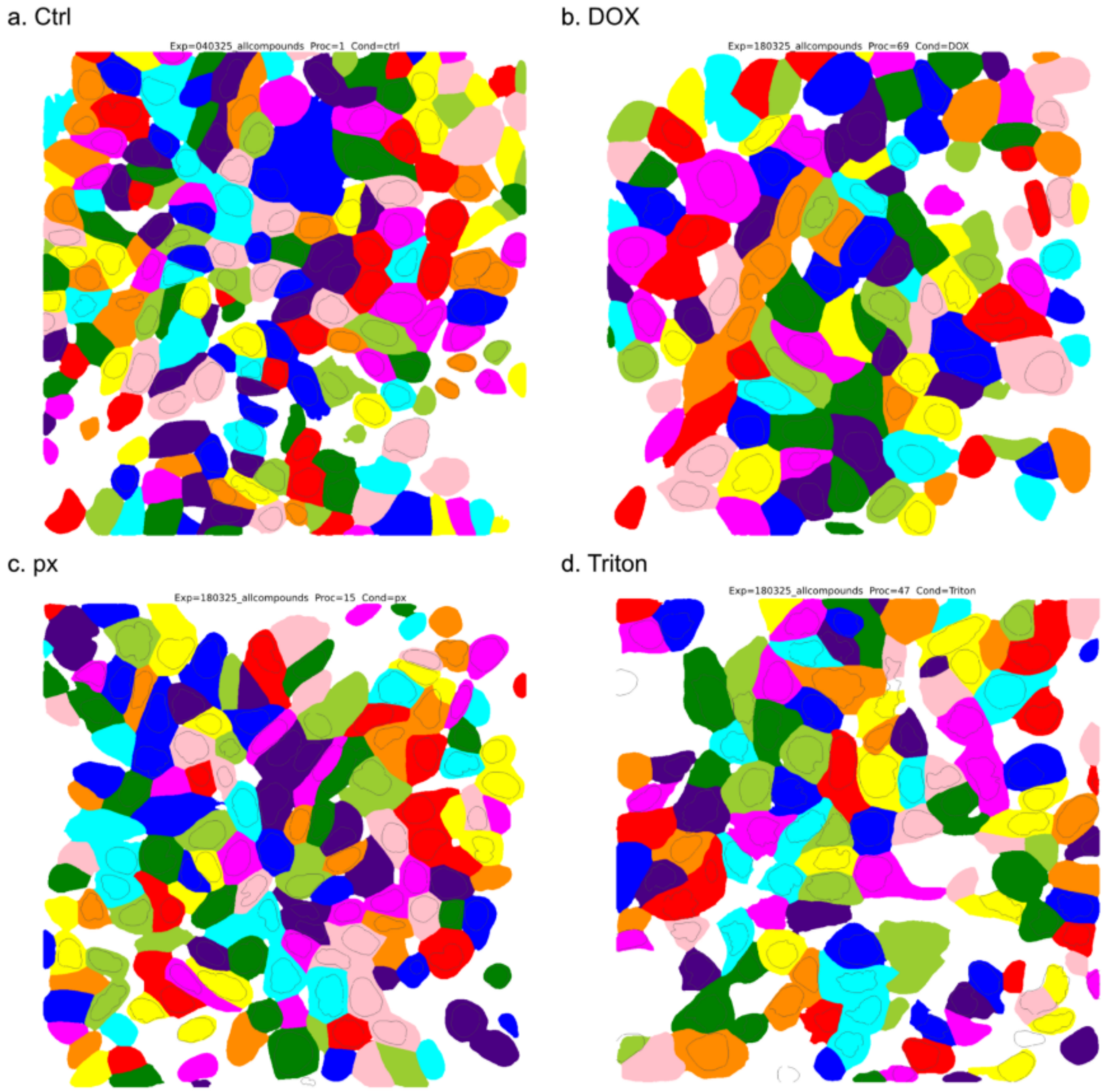


**Figure 9:** Representative raw actin (colored mask) and overlayed nucleus (non-colored masks) Caco-2 segmentations from Cellpose 3.0 for a) control (ctrl), b) doxorubicin (DOX), c) L-penetramax (px) and d) Triton™ X-100 (Triton).

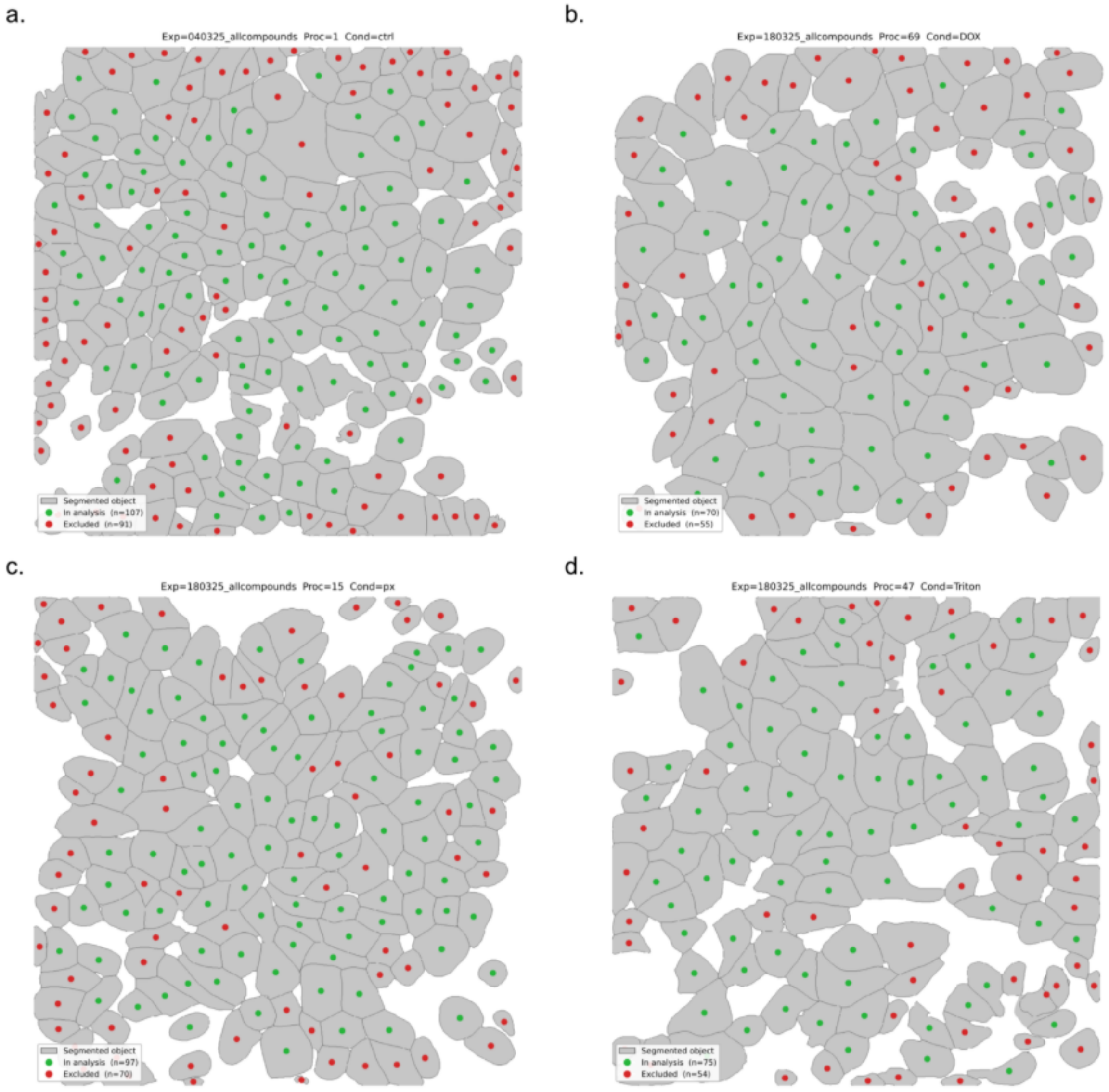


**Figure 10:** Representative Caco-2 final masks (green) consisting of a nucleus and actin paired and masks not included in analysis (red) for a) control (ctrl), b) doxorubicin (DOX), c) L-penetramax (px) and d) Triton™ X-100 (Triton).

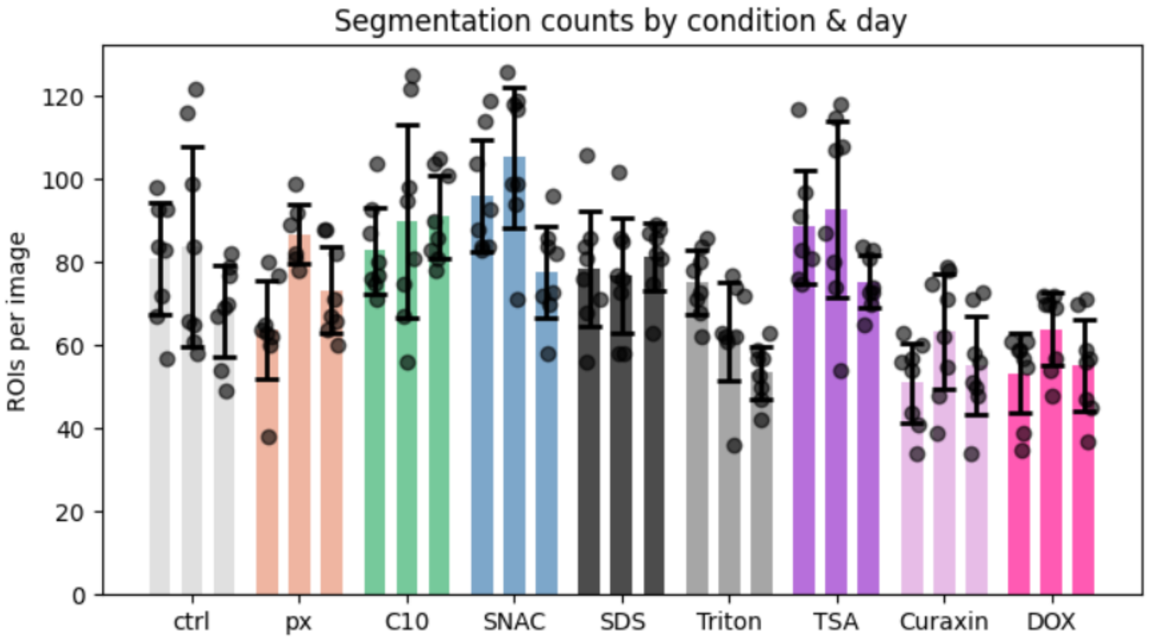


**Figure 11:** Segmentation counts per biological replicate for Caco-2 cells under exposure with hHBSS (ctrl), L-penetramax (px), sodium caprate (C10), sodium N-[8-(2-hydroxybenzoyl) amino] caprylate (SNAC), sodium dodecyl sulfate (SDS), Triton™ X-100 (Triton) for 30 min and 48h exposure with trichostatin A (TSA), curaxin CBL0137 (Curaxin) and doxorubicin (DOX) (i = 4, n = 2, N = 3).

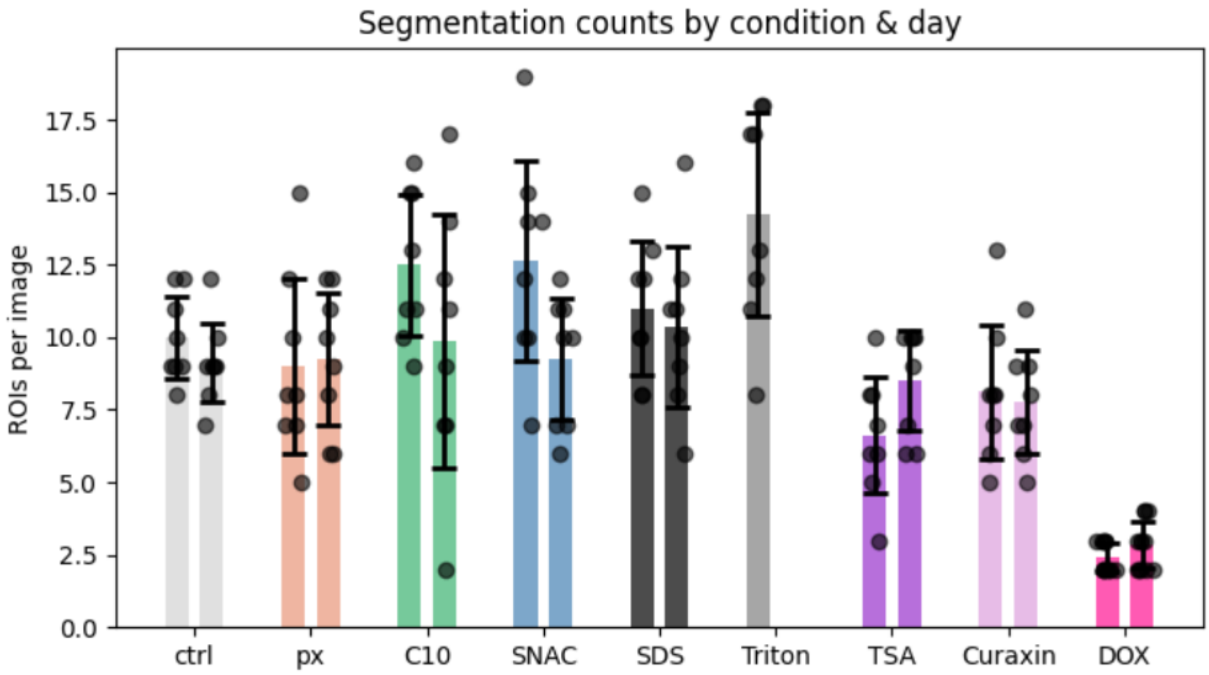


**Figure 12:** Segmentation counts per biological replicate for HeLa cells under exposure with hHBSS (ctrl), L-penetramax (px), sodium caprate (C10), sodium N-[8-(2-hydroxybenzoyl) amino] caprylate (SNAC), sodium dodecyl sulfate (SDS), Triton™ X-100 (Triton) for 30 min and 48h exposure with trichostatin A (TSA), Curaxin CBL0137 (Curaxin) and doxorubicin (DOX) (i = 4, n = 2, N = 2).

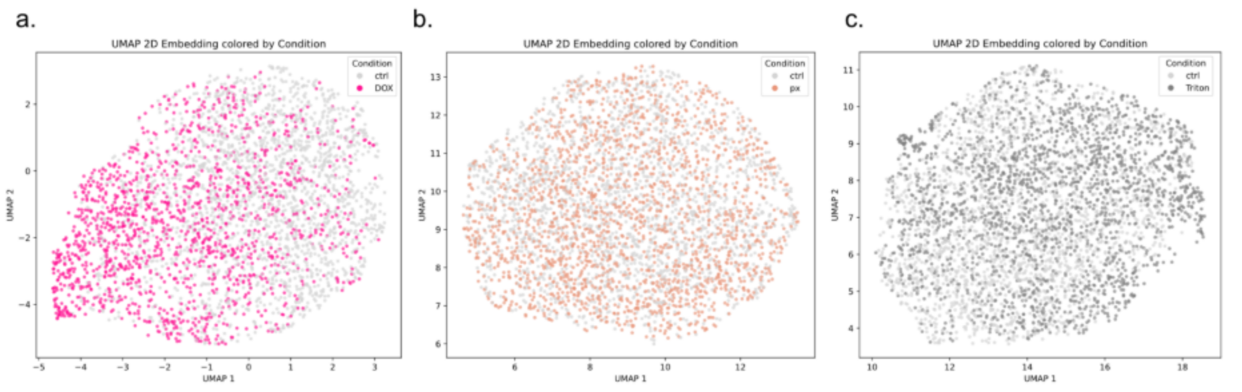


**Figure 13:** UMAP-embedding of the extracted features for control (ctrl) and treated Caco-2 monolayer cells for (a) control (ctrl, grey) and doxorubicin (DOX, pink), (b) control (ctrl, grey) and L-penetramax (px, orange) and (c) control (ctrl, grey) and Triton™ X-100 (Triton, dark grey).

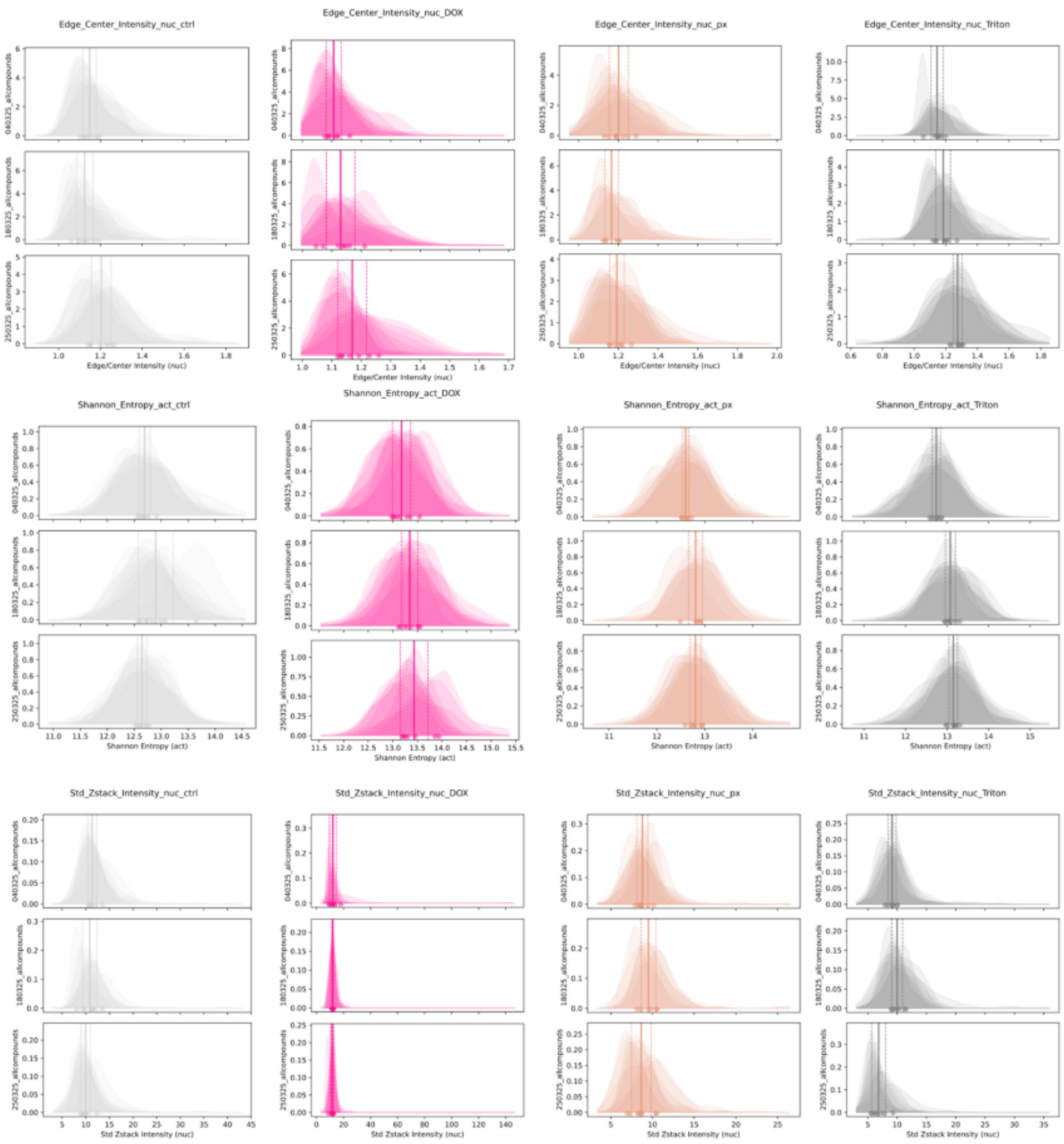


**Figure 14:** Feature distribution for three selected features for Caco-2 monolayer cells, plotted for each biological replicate separately. Data is shown for control (ctrl, grey), 30 min exposure with L-penetramax (px, orange) and Triton™ X-100 (triton, grey) and for 48h exposure with doxorubicin (DOX, pink). Each FOV is plotted as their own KDE. We see very little difference from biological replicate to replicate and very little heterogeneity across FOVs. Mean and std indicated as overlayed lines.

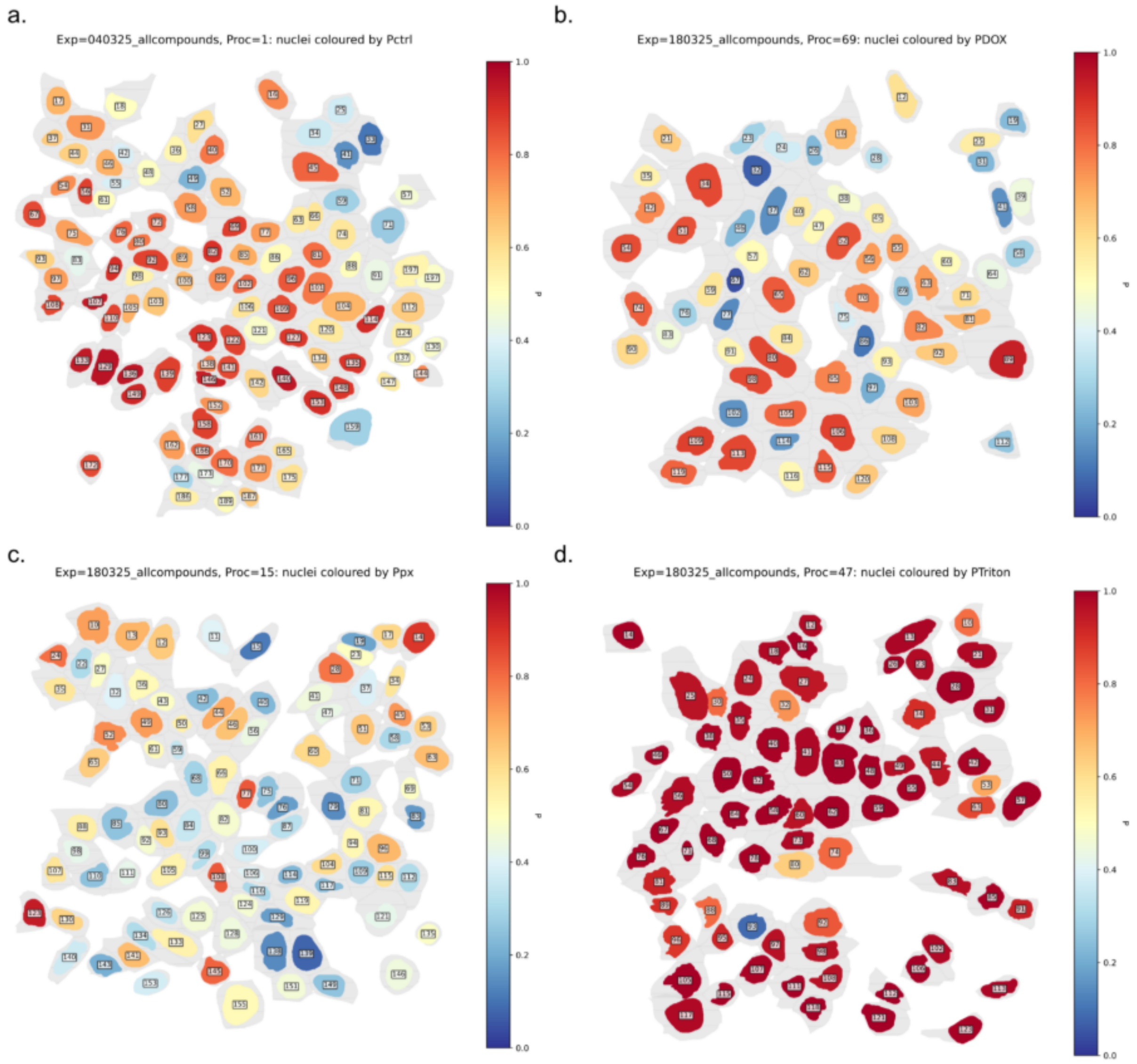


**Figure 15:** Representative FOV with segmented Caco-2 cells colored after XGBoost single-cell probability to belong to its own class for (a) control (ctrl), (b) doxorubicin (DOX), (c) L-penetramax (px) and (d) Triton™ X-100 (Triton) for a binary control (ctrl) vs. treatment classification

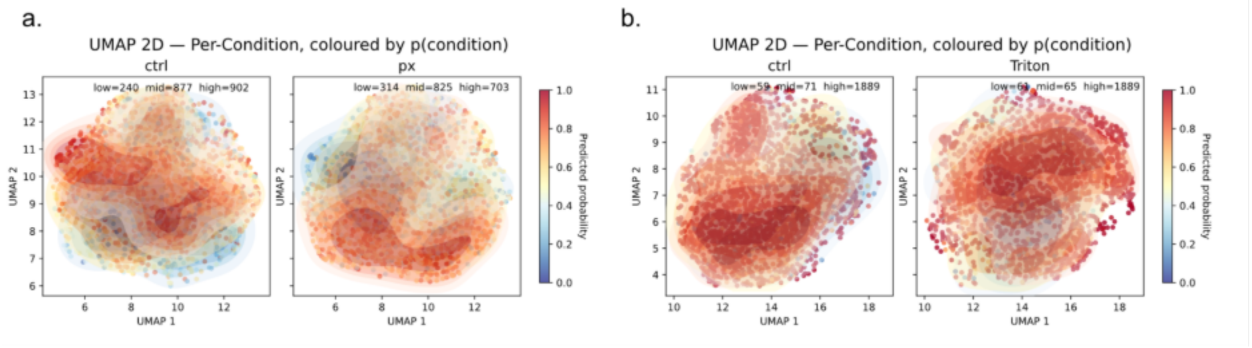

**Figure 16:** UMAP-embedding of feature-space for (a) control (ctrl) and L-penetramax-treated Caco-2 monolayer cells (px) and (b) control (ctrl) and Triton™ X-100-treated Caco-2 monolayer cells (Triton). The colorbar indicates single-cell probabilities.

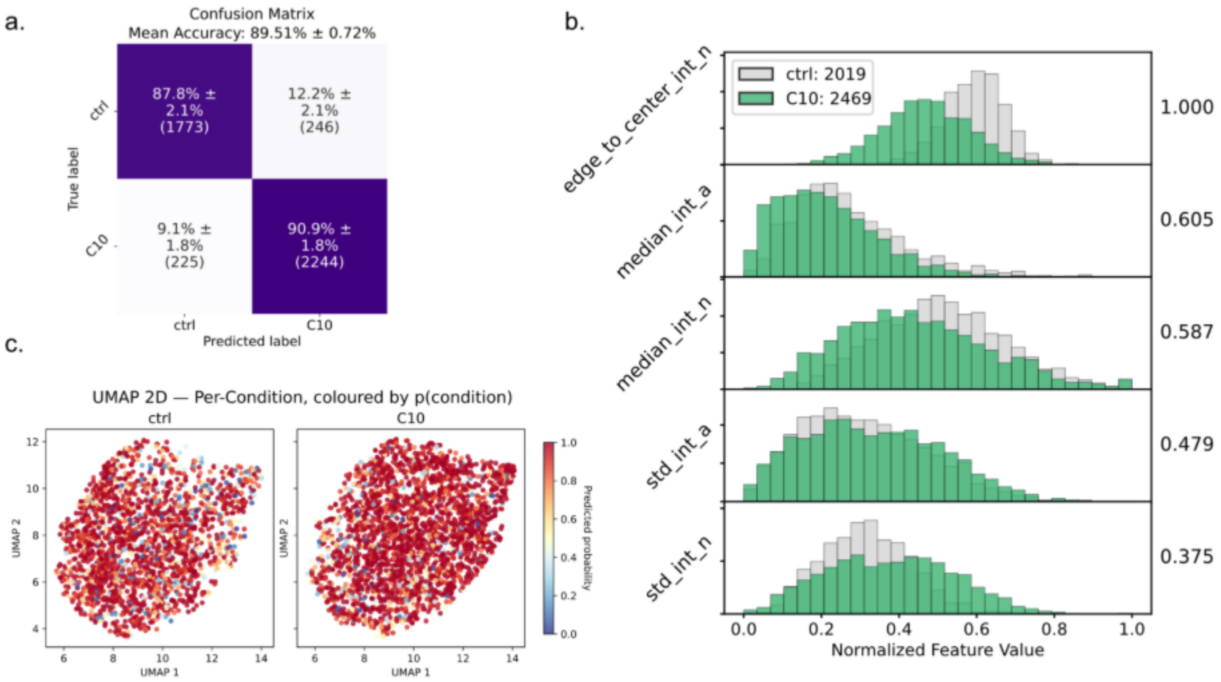


**Figure 17:** XGBoost classification of the binary control (ctrl, grey) vs. sodium caprate (C10, green) exposed Caco-2 cells. (a) confusion matrix for five-fold cross validation. (b) top 5 features by SHAP feature importance. (c) 2D UMAP of feature-space with single cell prediction probability as colorcode.

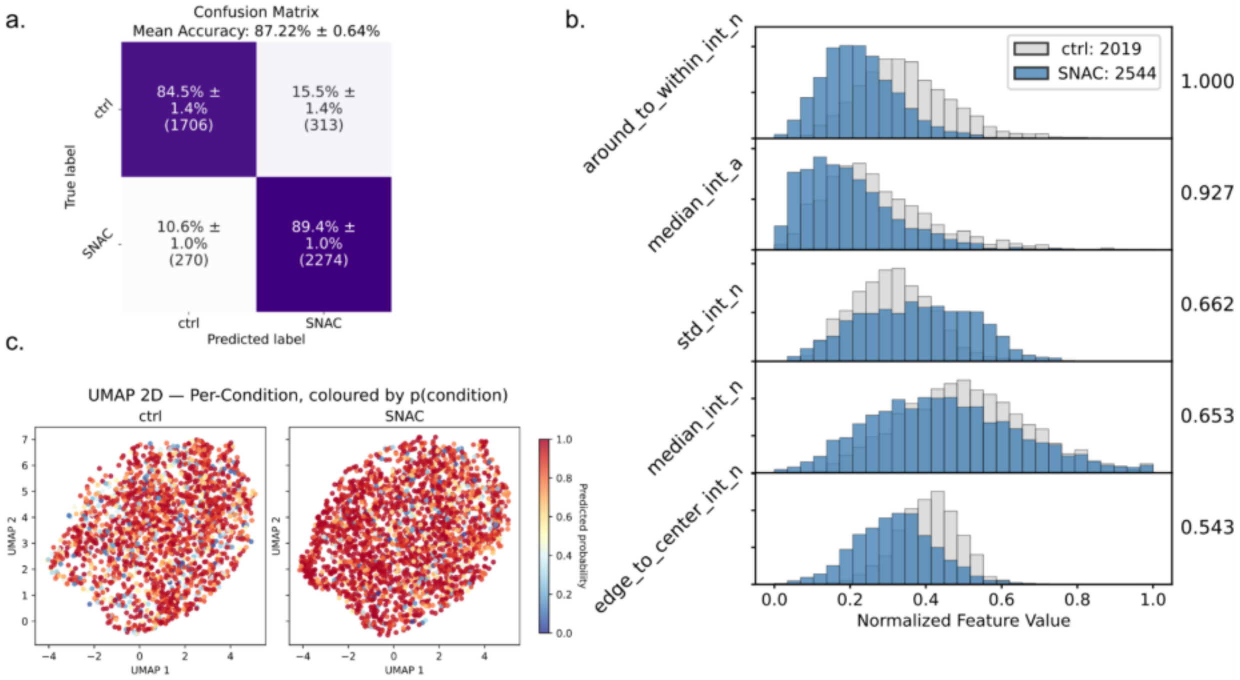


**Figure 18:** XGBoost classification of the binary control (ctrl, grey) vs. sodium N-[8-(2-hydroxybenzoyl) amino] (SNAC, blue) exposed Caco-2 cells. (a) confusion matrix for five-fold cross validation. (b) top 5 features by SHAP feature importance. (c) 2D UMAP of feature-space with single-cell prediction probability as colorcode.

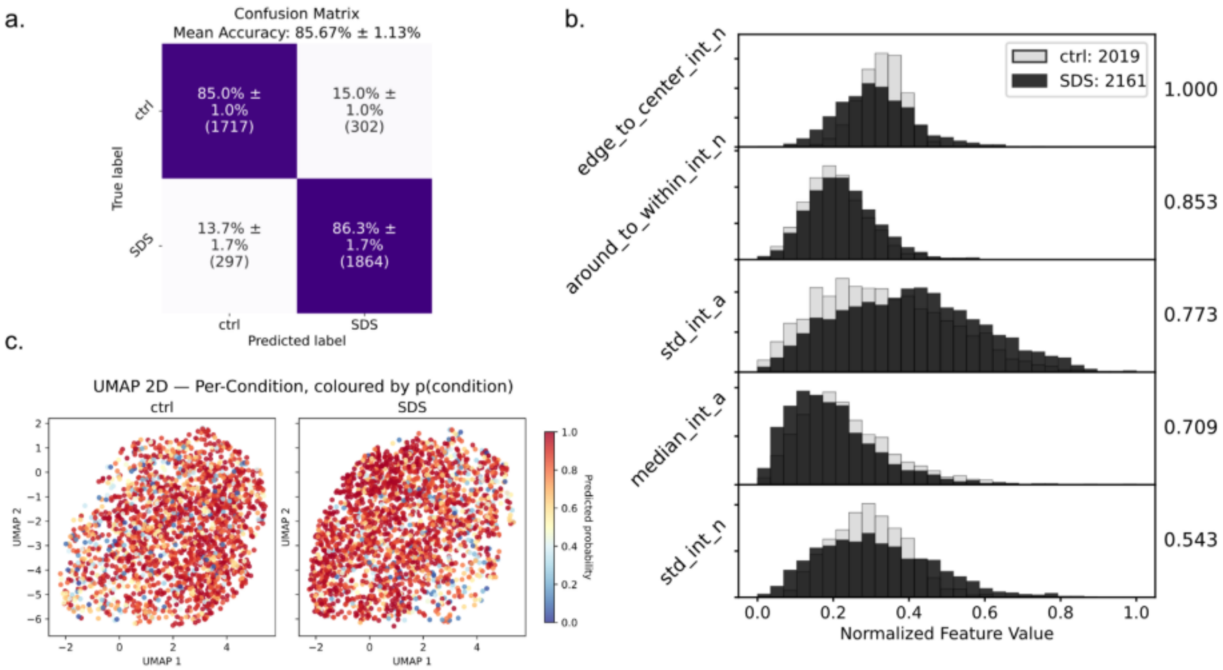


**Figure 19:** XGBoost classification of the binary control (ctrl, grey) vs. sodium dodecyl sulfate (SDS, black) exposed Caco-2 cells. (a) confusion matrix for five-fold cross validation. (b) top 5 features by SHAP feature importance. (c) 2D UMAP of feature-space with single-cell prediction probability as colorcode.

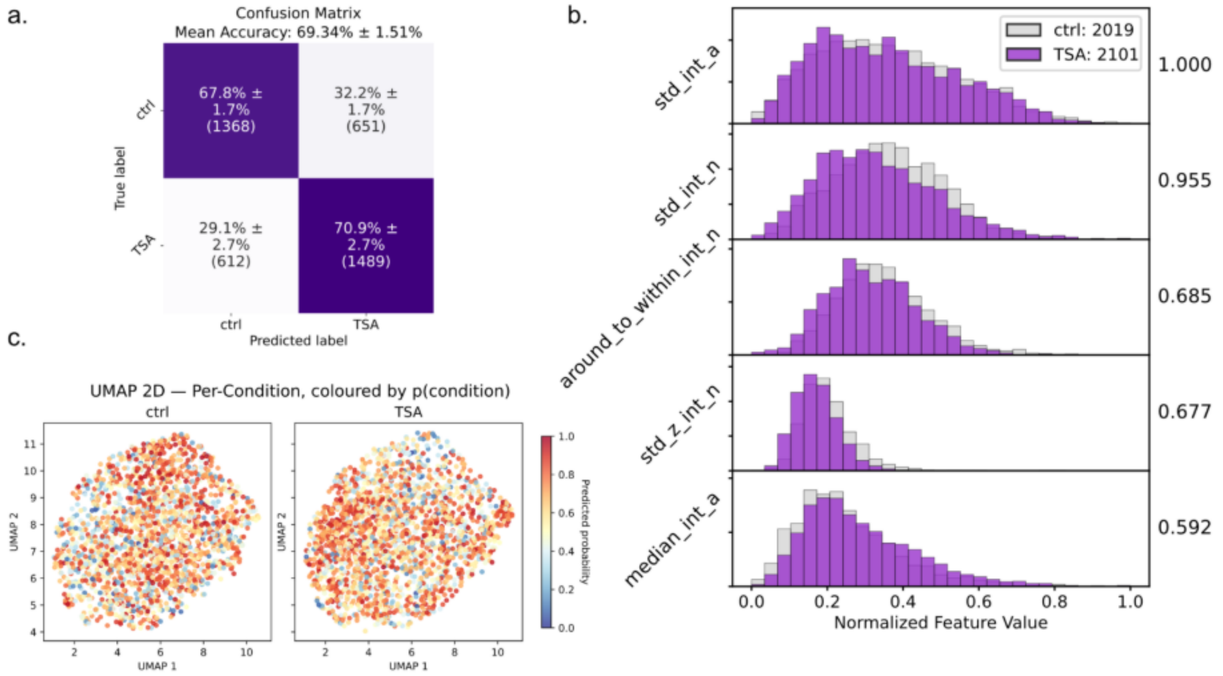


**Figure 20:** XGBoost classification of the binary control (ctrl, grey) vs. Trichistatin A (TSA, purple) exposed Caco-2 cells. (a) confusion matrix for five-fold cross validation. (b) top 5 features by SHAP feature importance. (c) 2D UMAP of feature-space with single-cell prediction probability as colorcode.

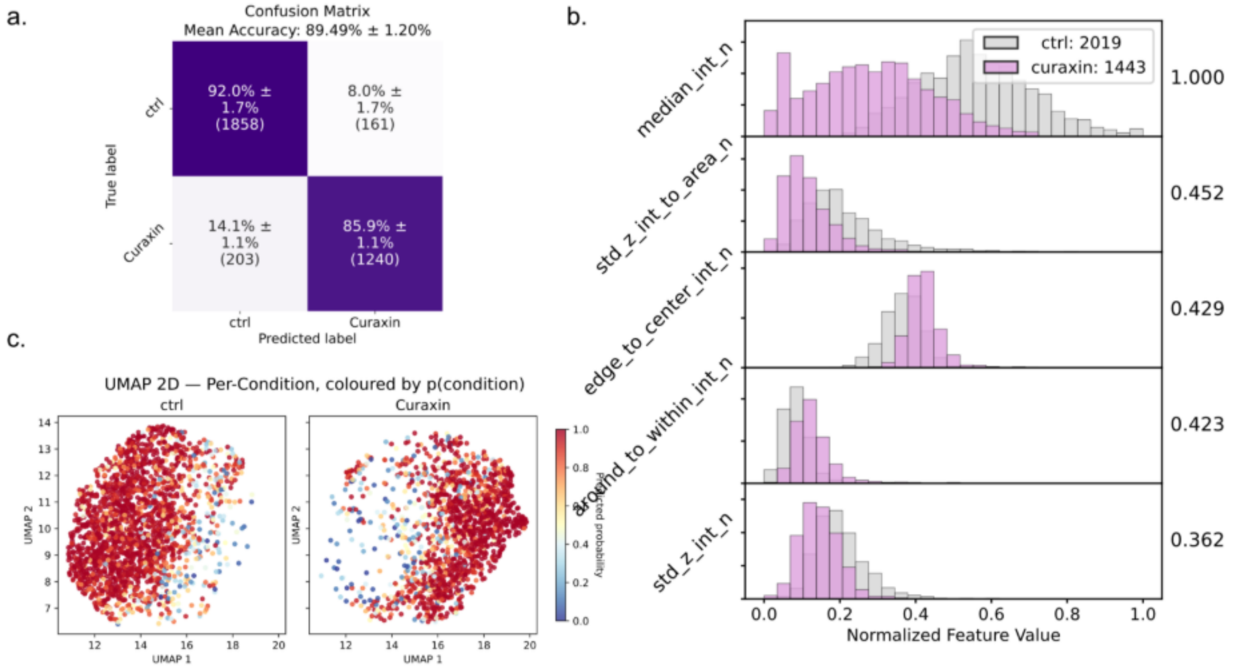

**Figure 21:** XGBoost classification of the binary control (ctrl, grey) vs.. Curaxin CBL0137 (Curaxin, light pink) exposed Caco-2 cells. (a) confusion matrix for five-fold cross validation. (b) top 5 features by SHAP feature importance. (c) 2D UMAP of feature-space with single-cell prediction probability as colorcode.

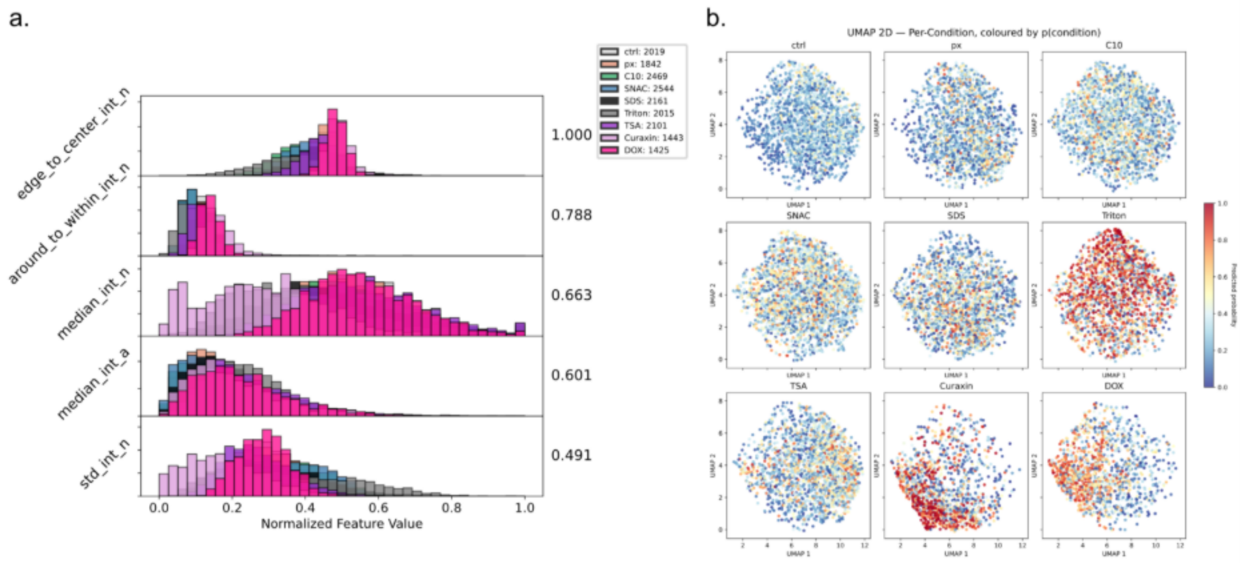


**Figure 22:** XGBoost 9-class classification for Caco-2 cells exposed with L-penetramax (px), sodium caprate (C10), sodium N-[8-(2-hydroxybenzoyl) amino] caprylate (SNAC), sodium dodecyl sulfate (SDS), Triton™ X-100 (Triton) for 30 min and 48h exposure with Trichostatin A (TSA), Curaxin CBL0137 (Curaxin) and doxorubicin (DOX). (a) top 5 features by SHAP feature importance. (b) 2D UMAP of feature-space with single-cell prediction probability as colorcode.

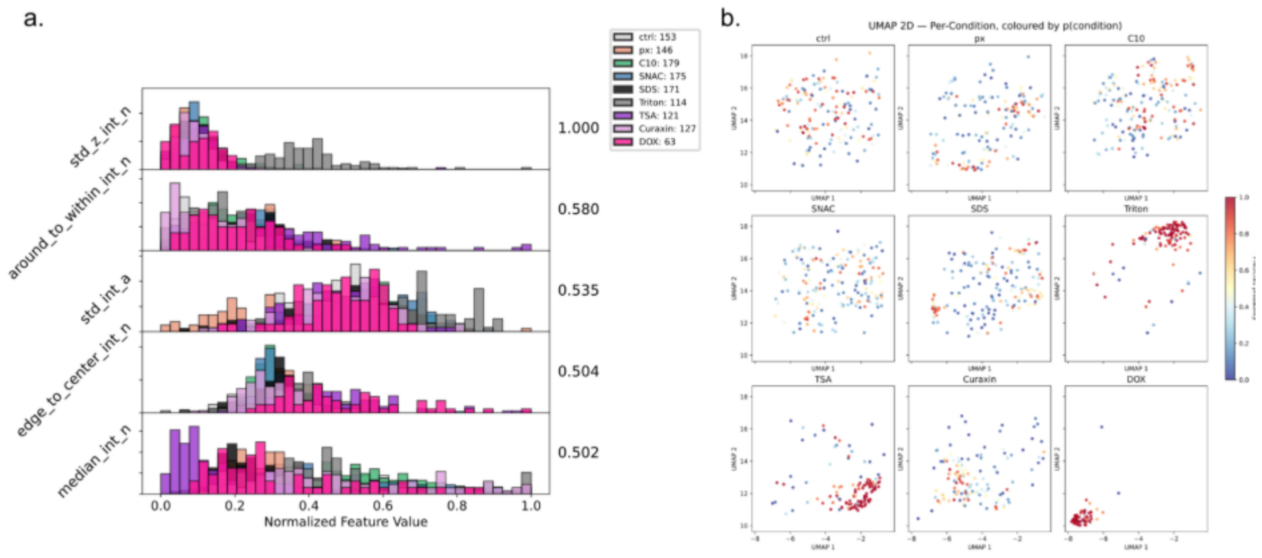


**Figure 23:** XGBoost 9-class classification for HeLa cells exposed with L-penetramax (px), sodium caprate (C10), sodium N-[8-(2-hydroxybenzoyl) amino] caprylate (SNAC), sodium dodecyl sulfate (SDS), Triton™ X-100 (Triton) for 30 min and 48h exposure with Trichostatin A (TSA), Curaxin CBL0137 (Curaxin) and doxorubicin (DOX). (a) top 5 features by SHAP feature importance. (b) 2D UMAP of feature-space with single-cell prediction probability as colorcode.

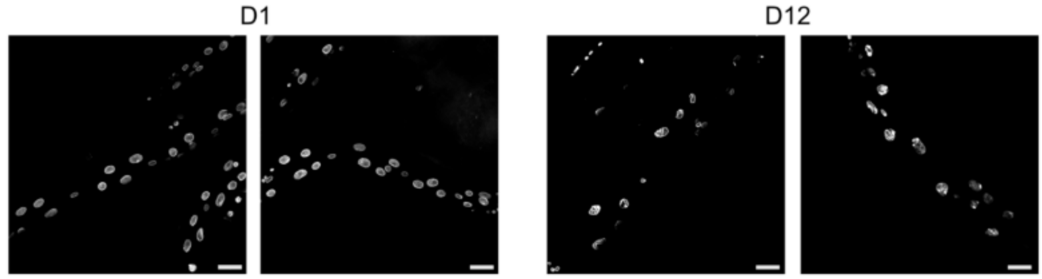


**Figure 24:** Representative microscopy images of nuclei of *C. elegans*, expressing a fluorescent reporter of the nuclear lamina protein lamin (LMN-1::GFP) *in vivo* for day 1 (young, D1) and day 12 (aged, D12) (scalebar = 20 µm, n = 1, N = 45-48).

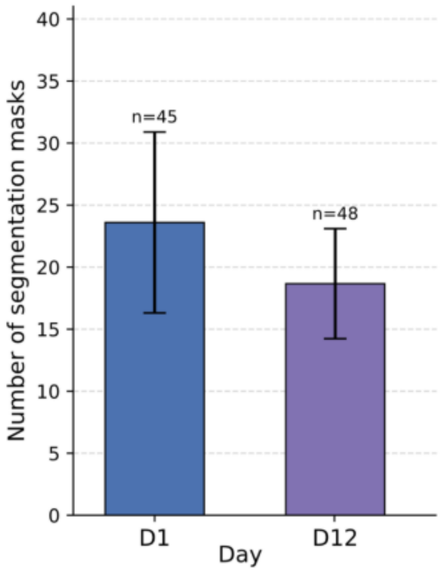


**Figure 25:** Segmentation count for *C. elegans* D1 and D12 in each acquired image (n = 1, N = 45-48).

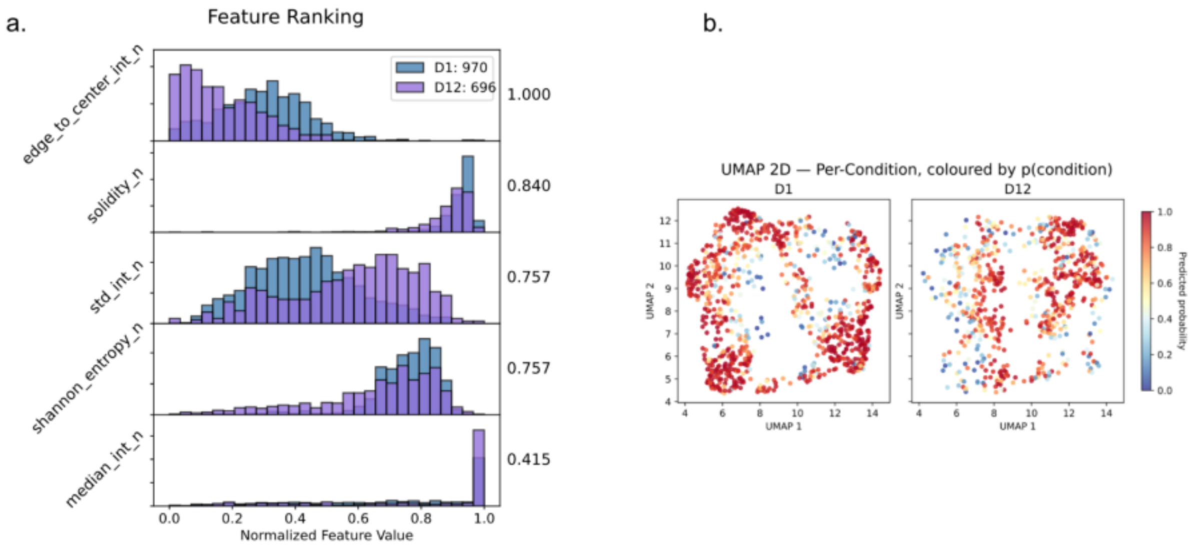


**Figure 26:** XGBoost classification of *C. elegans* D1 vs. D12. (a) top 5 features by SHAP feature importance. (b) 2D UMAP of feature-space with single-cell prediction probability as colorcode.
